# Fast and universal single molecule localization using multi-dimensional point spread functions

**DOI:** 10.1101/2023.10.17.562517

**Authors:** Mengfan Li, Wei Shi, Sheng Liu, Shuang Fu, Yue Fei, Lulu Zhou, Yiming Li

**Affiliations:** Department of Biomedical Engineering, Guangdong Provincial Key Laboratory of Advanced Biomaterials, Southern University of Science and Technology, Shenzhen, China; Department of Physics and Astronomy, University of New Mexico, Albuquerque, NM, USA

## Abstract

The recent development of single molecule imaging techniques has enabled not only high accuracy spatial resolution imaging but also information rich functional imaging. Abundant information of the single molecules can be encoded in its diffraction pattern and be extracted precisely (e.g. 3D position, wavelength, dipole orientation). However, sophisticated high dimensional point spread function (PSF) modeling and analyzing methods have greatly impeded the broad accessibility of these techniques. Here, we present a graphics processing unit (GPU)-based B-spline PSF modeling method which could flexibly model high dimensional PSFs with arbitrary shape without greatly increasing the model parameters. Our B-spline fitter achieves 100 times speed improvement and minimal uncertainty for each dimension, enabling efficient high dimensional single molecule analysis. We demonstrated, both in simulations and experiments, the universality and flexibility of our B-spline fitter to accurately extract the abundant information from different types of high dimensional single molecule data including multicolor PSF (3D + color), multi-channel four-dimensional 4Pi-PSF (3D + interference phase) and five-dimensional vortex PSF (3D + dipole orientation).

## Introduction

Single-molecule imaging is a powerful tool for investigating the spatial organization and functional information of molecular machines which underpin a myriad of biological processes. Especially, single molecule localization microscopy (SMLM) has overcome the diffraction limit and emerged as an important member of the super resolution microscopy methods. By sparsely activating single molecules in the wide-field, one can localize the spatial coordinates of each individual molecule with high resolution. As a result, fluorescent labeled biological structures can be imaged at resolutions close to the molecular scale (∼10 nm). Earlier efforts on SMLM focused on obtaining the super-resolved 3D positions from the shape of the diffraction limited single molecule images(*1*). In the past two decades, many other properties were also encoded to the single molecule diffraction pattern, such as color identity(*2-4*), fluorescence polarization(*5-8*), interference phase(*9-11*) etc. The resultant multi-dimensional super-resolution images enabled the construction of functional maps of varying new parameters at the super-resolution level(*12*).

Gaussian PSF model was the most widely used PSF model to extract the properties of the diffraction pattern (e.g., width, photons, locations) which were then converted to the 3D position(*13*). Although Gaussian PSF is computationally efficient, only limited PSF models can be analyzed by Gaussian PSF model. The recent development of spline PSF model was able to analyze experimental PSFs with arbitrary shape(*14-16*). Especially, if it is integrated with graphics processing unit, the spline PSF fitter achieved more than 100,000 fits/s, enabling real time single molecule localization(*15*). Furthermore, it can also be used to fit the single molecule data with a spline interpolation of an arbitrary analytical PSF model. Single molecule images contain rich information including both spatial and functional messages of individual molecules. To extract more information from such single molecules, specialized fitters are normally needed. By interpolating multi-dimensional PSF models, spline PSF fitter is ideal for extracting the rich information embedded in the single molecule data.

However, spline PSF fitter has so far been limited to 3D PSF model. One major obstacle to using the conventional spline PSF fitter for analyzing PSF models with higher dimension is the large number of spline coefficients needed, resulting in high memory demand. For a typical 3D PSF model by z scanning fluorescent beads with size of 31×31×100 pixels (∼3 µm×3 µm ×2 µm for pixel size of 100 nm in x, y and 20 nm in z), 64×31×31×100 ≈ 6.1×10^6^ cubic spline coefficients (23.4 MB for float data type) are needed. If we add another dimension with 50 sampling points, 256×31×31×100×50 ≈ 12.3×10^8^ coefficients (4.5 GB for float data type) will be needed. The number of coefficients increases by two orders of magnitude when the PSF model is increased by one dimension. It is therefore difficult to apply conventional spline interpolation for high dimensional single molecule data. Furthermore, the speed of the data analysis decreases dramatically with the increase of the dimension in the fitting PSF model. A fast data analysis pipeline is also required.

Here, we developed a B-spline interpolated PSF model that can be represented by linear combination of basis functions. For B-spline interpolation, there is only one B-spline coefficient associated with each grid point of the data interpolated(*17*). It therefore greatly reduces the order of magnitude of the spline coefficients that is needed to describe a multi-dimensional PSF model. As a result, cubic B-spline coefficients with 24 MB memory size is enough to describe a model with sampling data point of 31×31×100×50, while 4.5GB memory size is needed for conventional cubic spline coefficients. To speed up the data analysis process with B-spline interpolated PSF model, we implemented the multi-dimensional PSF fitting with GPU. To demonstrate the wide applicability of the new B-spline based multi-dimensional PSF fitter, we used the new fitter to extract multi-dimensional parameters from multicolor PSF (3D + color), multi-channel four-dimensional 4Pi-PSF (3D + interference phase) and five-dimensional vortex PSF (3D + dipole orientation).

## Results

### Interpolation of multi-dimensional PSFs using B-spline

The PSF is the impulse response of an imaging system which reflects the properties of the image formation process. It is often recorded in a 2D image. Many physical parameters (e.g., locations(*1*), emission wavelength(*2, 3*), fluorescence dipole orientation(*7*)) of the single point emitter could change the shape of the PSF (**Fig. 1a**). Therefore, by fitting the 2D single molecule images with a proper PSF model, one can precisely extract the physical properties of single emitters. In the past decade, more and more engineered PSFs were introduced, allowing for better 3D localization(*18-20*), color separation(*21*) and orientation estimation(*7, 8*). However, the analysis methods for different types of single molecule patterns are often sophisticated and specialized software is needed, impeding the broad application of multi-dimension functional imaging of single molecules.

**Fig. 1.**
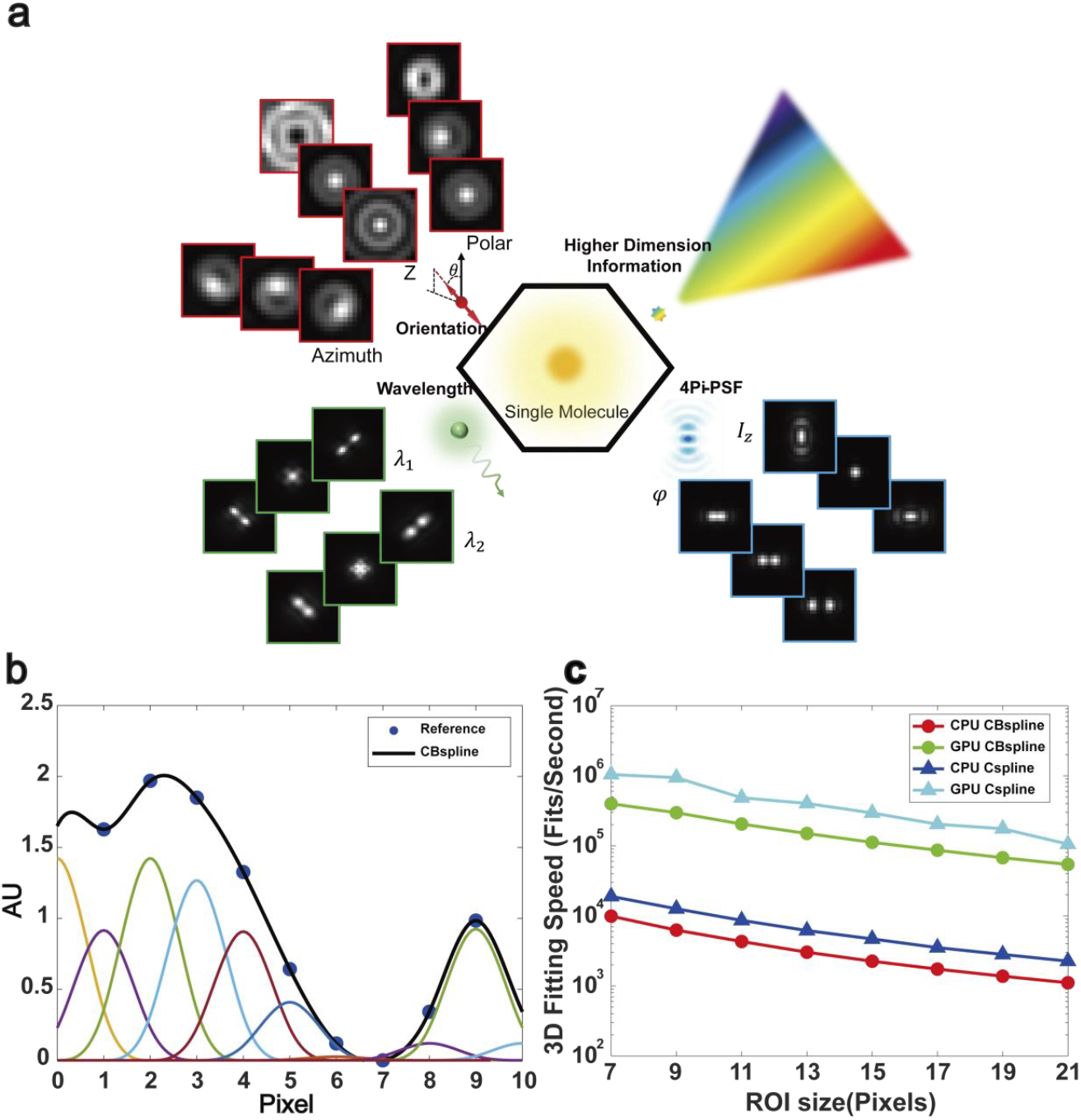
Schematic diagram of B-spline based multi-dimensional PSF model. **a**, Single molecule images contain multi-dimensional information. **b**, CBspline interpolates discrete points with cubic curves calculated from the weighted sum of the basis functions. **c**, Comparation of the speed of Cspline (standard cubic spline) and CBspline (cubic B-spline) fit.

Splines are piecewise polynomials that provide a simple mechanism to convert discrete signals into continuous ones. They have been extensively used in surface approximation and multi-dimensional data fitting. B-spline is one type of spline that is constructed as a linear combination of basis functions. The B-splines can be made to give the same result as standard splines, within numerical rounding error (**Supplementary Fig. 1**). There are several advantages of B-splines compared to the standard splines: 1) The number of B-spline coefficients needed per mesh point does not increase with dimensionality. In contrast, conventional splines required 4^d^ coefficients per mesh point for cubic splines, where d is the number of dimensions. Therefore, a factor of 64 in storage is saved for 3D PSF when using cubic B-spline instead of standard cubic spline. For higher dimensional PSFs, the difference in storage requirements will be more significant; 2) Using different basis functions, B-spline is more convenient for signal processing such as signal differentiation, filtering, smoothing, and least square approximations(*17*). Especially for the noisy PSF models measured experimentally, a certain degree of smoothness is needed for PSF modeling to avoid artifacts(*15*); 3) The change of the local control points does not globally change the shape of the whole curve which makes the model interpolated more stable. **Fig. 1b** shows the example of the construction of a one-dimensional B-spline. The interpolated values are the summation of the basis functions with different amplitudes given by the B-spline coefficient at each grid point. Setting the magnitude of each basis function reduces the size of the parameters of B-spline compared to that of individual expression of each line segment. Therefore, it is more convenient for building a higher dimensional PSF as the number of coefficients for each grid point does not increase with the dimensions.

In our previous work, we used the standard cubic spline (Cspline) to realize 3D fitting and used GPU parallel computing to accelerate the fitting speed(*15*). In this work, we also implemented cubic B-spline (CBspline) based multi-dimensional PSF fitting in GPU and improved the speed almost two orders of magnitude compared to the CPU based calculation (**Fig. 1c**). Our GPU based CBspline fitter achieved Cramer-Rao-Lower-Bound (CRLB) in all dimensions and in multi-channel PSF fitting (**Supplementary Figs. 2 and 3, Supplementary note 1 and 2**). Especially for the 4D and 5D PSF model, the memory needed for the Cspline coefficients is in the order of gigabytes and hundred gigabytes, while it only needs megabytes and hundred megabytes for CBspline coefficients (**Supplementary Table 1a**). Considering that the memory for a typical GPU is in the order of gigabytes, it is difficult to implement higher dimensional PSF fitting in GPU using Cspline. In terms of algorithms complexity, B-spline based PSF modeling has more multiplication and addition operations than standard spline based PSF modeling. In the 3D PSF case when it is still possible to store Cspline coefficients in a typical GPU, CBspline based PSF modeling is about 2.5 times slower compared to Cspline without considering the memory copying between host CPU and GPUs (**Fig. 1c**).

### Simultaneous measurement of 3D positions and color information using B-spline

To measure the color information of single molecules, approaches such as ratio metric multicolor(*22-24*), spectrometer(*25*) and PSF engineering(*3, 4*) were employed. However, the shape of PSF itself was normally encoded with wavelength information. From the perspective of Fourier optics, the wavelengths represent the spatial frequencies on the Fourier plane. For an objective with a fixed numerical aperture (NA) and collected photons with wavelength λ, the maximum frequency of the PSF is 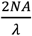 . Thus, PSF with longer wavelength has a smaller cutoff frequency and a slightly larger shape (**Fig. 2f, Supplementary Fig. 4**). Furthermore, optical aberrations are also different for different wavelengths due to dispersion. Therefore, there is normally a subtle difference among PSFs with different wavelengths. Recently, neural networks were used to extract the color information from unmodified microscopes(*2*). However, it is not clear whether the achieved localization accuracy and color separation accuracy achieved the theoretical optimal performance. Moreover, proper training of a neural network is still challenging for different experimental conditions.

**Fig. 2.**
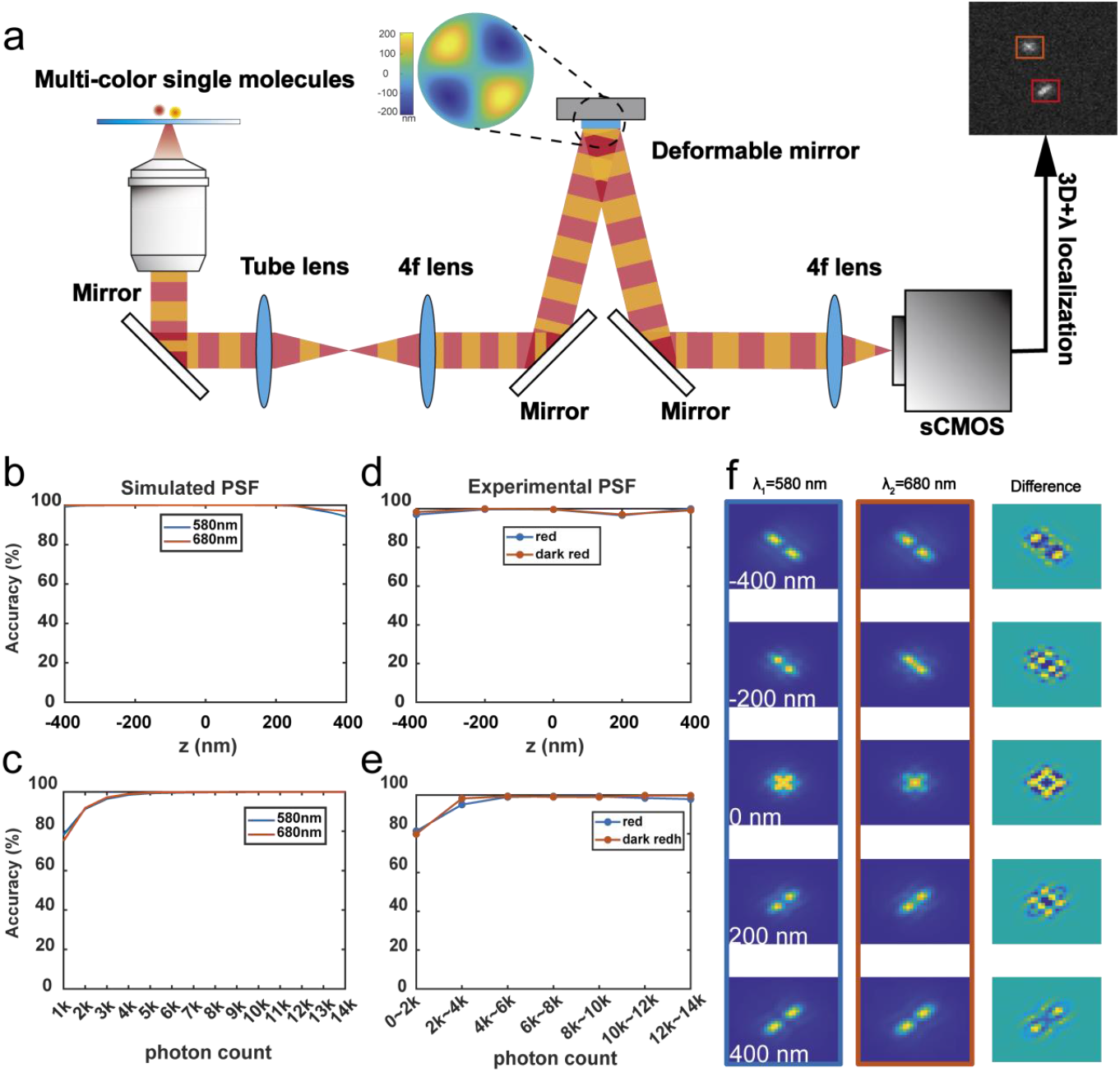
Simultaneous measurement of 3D positions and color information using CBspline. **a**, Schematic diagram of the optical path for DMO PSF engineering at different wavelengths using a deformable mirror. Fluorescence of different emission wavelengths were passed through the same DM. **b** and **c** are the color classification accuracy as function of different axial positions (single molecules with 5,000 photons and 20 background photons) and different photon count using simulated data, respectively. **d** and **e** are the same **b** and **c**, but using experimental PSF acquired from red and dark red fluorescent beads (emission peak at 605 nm and 680 nm, respectively). The photon distribution of the beads evaluated can be found in **Supplementary Fig. 7**. Only beads with more than 4,000 photons were used for evaluation. **f**, Simulated DMO PSFs with different emission wavelengths (580 nm and 680 nm) and their difference. The simulated data are derived from the vectorial PSF model and include refractive index mismatch induced aberrations (**Methods**).

Here, we built a multi-color 3D PSF model both theoretically and experimentally (**Methods**). We used CBspline to interpolate the 3D+λ PSF model and fit the single molecule data with the CBspline interpolated PSF model. We first evaluated the performance of the widely used astigmatic PSF model (80 nm astigmatism) for color separation and 3D localization precision. PSFs of two different wavelengths (580 nm and 680 nm) were used. We systematically evaluate the color separation of these two PSFs at different z positions and signal to noise ratio (SNR). As shown in **Supplementary Fig. 5a**, PSFs at different z positions show different color separation accuracy. We then quantify the color separation accuracy for photon number from 1,000 to 14, 000. Axial range of 400 nm around the focus was used for evaluation. For a typical single molecule emission (2,000 photons and 20 background photons per pixel), the color separation accuracy is about 80.1% for PSFs at 580 nm, while it is 77.5% for PSFs at 680 nm (**Supplementary Fig. 5b**). When the photon number is increased to 10, 000, the color separation accuracy achieved about 97% for both PSFs.

Since our B-spline fitter is applicable to PSFs with arbitrary shape, we also constructed deformable mirror based optimal (DMO) PSF of different wavelengths, which we showed could achieve optimal 3D localization precision using deformable mirror(*20*) (**Fig. 2a**). DMO PSF with 1.2 µm depth of field was used in this work. DMO PSF showed not only superior performance on 3D localization precision (**Supplementary Fig. 6**), but also much better color classification ability (**Figs. 2b** and **c, Supplementary Fig. 5**). The averaged axial 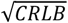 of DMO PSF is 21.1 nm while it is 31.1 nm for astigmatic PSF (2,000 signal photons and 20 background photons, **Supplementary Fig. 6**). The average color classification accuracy is more than 99.3% for molecules at z positions within 400 nm around the focus (**Fig. 2b**). As shown in **Fig. 2f**, the wavelength affects both the lobes’ distance and size, which results in good color separation accuracy. These results indicate that DMO PSF rapidly changes with the change of both wavelength and axial position.

We then evaluated the color separation ability of DMO PSF with experimental data. 40 nm diameter fluorescent beads with different emission wavelengths were used (605 nm and 680 nm, **Methods**). To quantify the color separation accuracy at different SNRs and axial positions, we acquired z bead stacks under different laser excitation power (**Methods**). For beads at different z positions (photon count range in 4,000 to 14,000, **Supplementary Fig. 7**), the averaged color distinction accuracy also exceeded 95% and reached 99.6% at focus position (**Fig. 2d**). The average accuracy of color classification exceeds 90% for beads with photons higher than 2,000 and 98% for beads with photons higher than 5,000 at the axial positions between -400 nm to 400 nm (**Fig. 2e**).

We then applied the 3D+λ fitter in the real biological experiments. Nuclear pore complex protein Nup96 and the mitochondrial outer membrane protein Tomm20 were individually labeled with AF647 and CF568. These labels have peak emission wavelengths of 665 nm and 583 nm, respectively (**Methods**). The samples were excited with 561 nm and 640 nm laser simultaneously. By adjusting the excitation power of these two lasers, the two dyes have roughly the same number of photons. The emitted fluorescence of these two dyes passed through the same imaging optical path and filter, and were recorded in the same area of the camera. The averaged photon counts/switch event for CF568 and AF647 were 5,264 and 9,393, respectively (**Supplementary Fig. 8**). As shown in **Fig. 3**, the mitochondria and nuclear pore complex (NPC) were very well distinguished. The octuple ring and two-layer morphology of the Nup96 residing NPC was also very nicely resolved, indicating that both good 3D localization and color separation accuracy were achieved by our B-spline fitter.

**Fig. 3.**
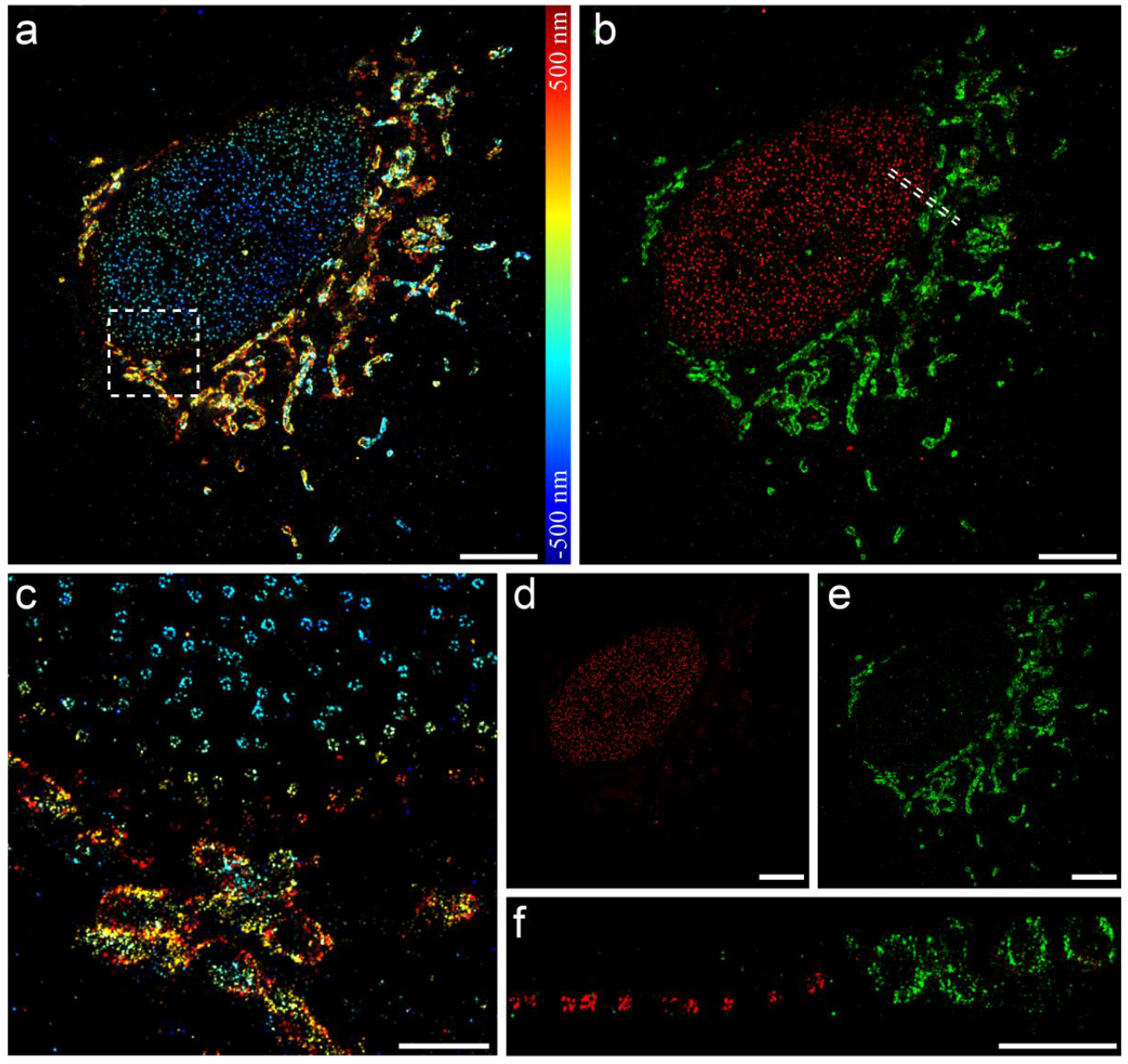
Simultaneous 3D imaging of nuclear pore and mitochondria using DMO PSF analyzed by CBsplines. **a**, 3D super resolution images of nuclear pore complex protein Nup96 and mitochondria outer membrane protein Tomm20. **b**, Color separation of Nup96 and Tomm20. **c**, Zoomed image of the boxed region in **a. d** and **e** are individual color images of **b. f**, Side view of boxed region in **b**. Scale bars, 5 μm (**a, b, d, e**), 1 μm (**c, f**).

### Analysis of multi-channel four dimensional 4Pi-PSF using B-spline

4Pi-SMLM uses two opposing objectives to coherently detect the single molecule fluorescence and achieved sub-10 nm isotropic 3D resolution even for single molecules with 250 photons collected by each objective(*26*). In 4Pi-SMLM, the change of axial position of the emitter will change the optical path length difference (OPD) within the interference cavity, thus changing the interference phase of the self-interfered individual photons. However, the OPD does not only depend on the axial position, but also many other disturbances (e.g., temperature change, inhomogeneous refractive index) during real experiments (**Supplementary Fig. 9a**). Furthermore, 3 or 4 interference phase channels are normally collected simultaneously to avoid axial regions where the interference intensity changes slowly with z(*27-29*). Therefore, 4Pi-PSF is intrinsic a multi-channel 4D PSF model (**Supplementary Fig. 9b**). To analyze 4Pi-SMLM data, different algorithms have been proposed(*9, 11, 27*). However, all of them needed specialized PSF fitter for data analysis. It often needs to derive the partial derivatives of the PSF models with respect to each dimension which is often not easy to be generalized.

Here, we developed a CBspline based multi-channel 4D PSF fitter which could directly interpolate the multi-channel 4Pi-PSF by CBspline and perform 4D localization (x, y, z, and phase) on 4Pi single molecule data. Similar to our previously published globLoc fitter(*30*), it is flexible to link or unlink parameters between channels. By globally fitting data from all phase channels, our CBspline fitter achieved CRLB in all dimensions (**Supplementary Fig. 3**). Furthermore, we implemented the multi-channel CBspline fitter in GPU and achieved around 60 times faster compared to CPU code (**Supplementary Fig. 10a**). We also demonstrated the ultra-high accuracy structure with 4Pi-SMLM imaging of Nup96 analyzed by our CBspline fitter (**Fig. 4a**). We fitted the line profile of the double layer intensity along z with double Gaussian function. The sigma of both Gaussian is below 10 nm, indicating sub-10 nm axial resolution achieved with 4Pi-SMLM imaging (**Figs. 4d-f**). Compared to our previous work using IAB-4Pi-PSF model and interpolating I, A, B matrices with 3D cubic splines(*11*), we used CBspline to directly interpolate the 4D PSF model. Although only 4Pi-PSF model was demonstrated in this work, B-spline fitter could be applied to other multi-channel 4D data without further derivation of model gradients.

**Fig. 4.**
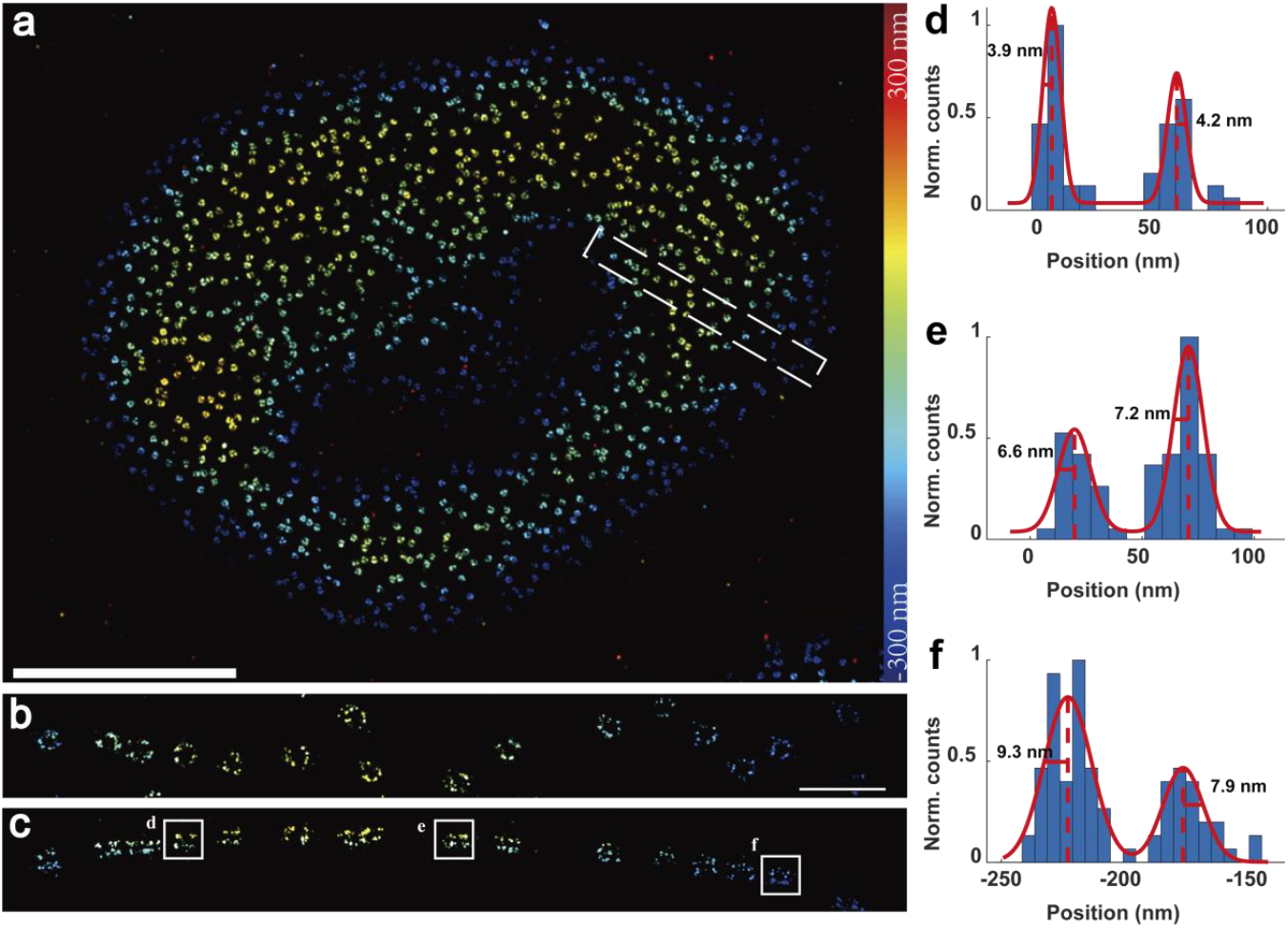
Multi-channel 4Pi-SMLM data analyzed by CBsplines. **a**, Top view of 4Pi-SMLM images of the nuclear pore complex protein Nup96. **b**, Zoomed image of the dash boxed area in **a. c** Side view of **b. d, e** and **f** are the axial intensity profiles of the regions as indicated in **c**. Double Gaussian was then fitted to these intensity profiles. Scale bars, 5 μm (**a)**, 500 nm (**b, c**).

### Simultaneous measurement of 3D positions and orientation information using B-spline

Fluorescence emission is well modeled by an oscillating electric dipole. The shape of the single molecule dipole imaged by the microscope is highly dependent on the orientation of the dipole. Especially for dipoles which are rotationally constrained in space, symmetric PSF models such as Gaussian would lead to localization bias(*31*). To extract the orientation information from the single molecule dipole emission pattern, complex vectorial PSF model based fitter are normally needed to account for the polarization effect. Many physical parameters are needed to build this model which are often difficult for non-optics experts. Due to the sophisticated calculation during the PSF computation, the fitting speed is normally in the range of tens fits/s, heavily limiting the imaging throughput of fluorescence anisotropy measurement. Since the accuracy of the orientation estimation is poor using the unmodified PSFs, we employed the recently published Vortex PSF to extract the emitters orientation(*7*). To localize the orientation of a dipole, three parameters are estimated, the polar angle respect to the optic axis *θ*, in plane azimuthal angle *ϕ* and the degree of oscillation *g*_2_ (a parameter quantifying the degree of rotational mobility). The final PSF is a weighted sum of freely rotating dipole PSF and fixed dipole PSF.

To verify the accuracy of five-dimensional CBspline fitter, we simulated a series of Vortex PSFs under various orientations and positions using vectorial PSF model. The fitting parameters include 3D coordinates *x, y, z*, photon count of the emitter *I*, the background photons per pixels *b*, orientation of the dipole *ϕ,θ* and the weight parameter (degree of orientational constraint) *g*_2_. The CRLB of all parameters were calculated by using the first derivative of PSF along each parameter derived by B-spline. The fitting precision evaluation was performed on molecules with 4,000 photons and 10 background photons. Here, we initialized the fitter with two different axial positions to prevent the fitter being trapped in local minimum. As shown in **Supplementary Fig**.**11**, the precision of all dimensions could reach the CRLB. Our GPU based 5D CBspline fitter improved computing speed by around 50 times compared to CPU based computing (**Supplementary Fig. 10b**).

We then applied our 5D CBspline fitter to analyze a previously published imaging data acquired from λ-DNA(*7*). As shown in the reconstructed super resolution image (**Fig. 5**), most of the molecules have an azimuthal angle perpendicular to the in plane orientation of λ-DNA (**Fig. 5a**). For the reconstructed 3D images as shown in **Fig. 5b**, the returned axial position could also show the intentionally tilted samples, which could be accurately resolved under different dipole orientation conditions. We then analyzed in detail the relative azimuthal angle with respect to the DNA axis. As shown in **Fig. 5c**, a single strand shows a mean azimuthal angle difference between the fluorophore and the DNA axis (red solid line) is 85.7° which is close to that found previously(*7*). We also analyzed the *g*_2_ on the same region of **Fig. 5c** and the distribution of *g*_2_ is peaked at 0.81 (**Fig. 5d**) which also agrees with previous analysis(*7*). In the zoomed *g*_2_ image as shown in **Fig. 5e**, *g*_2_ is about 0.76 in the single DNA strand (red arrow) while it is about 0.29 in the patches (green arrow). It indicates that the DNA might be partially detached or non-specifically bound at the patch locations.

**Fig. 5.**
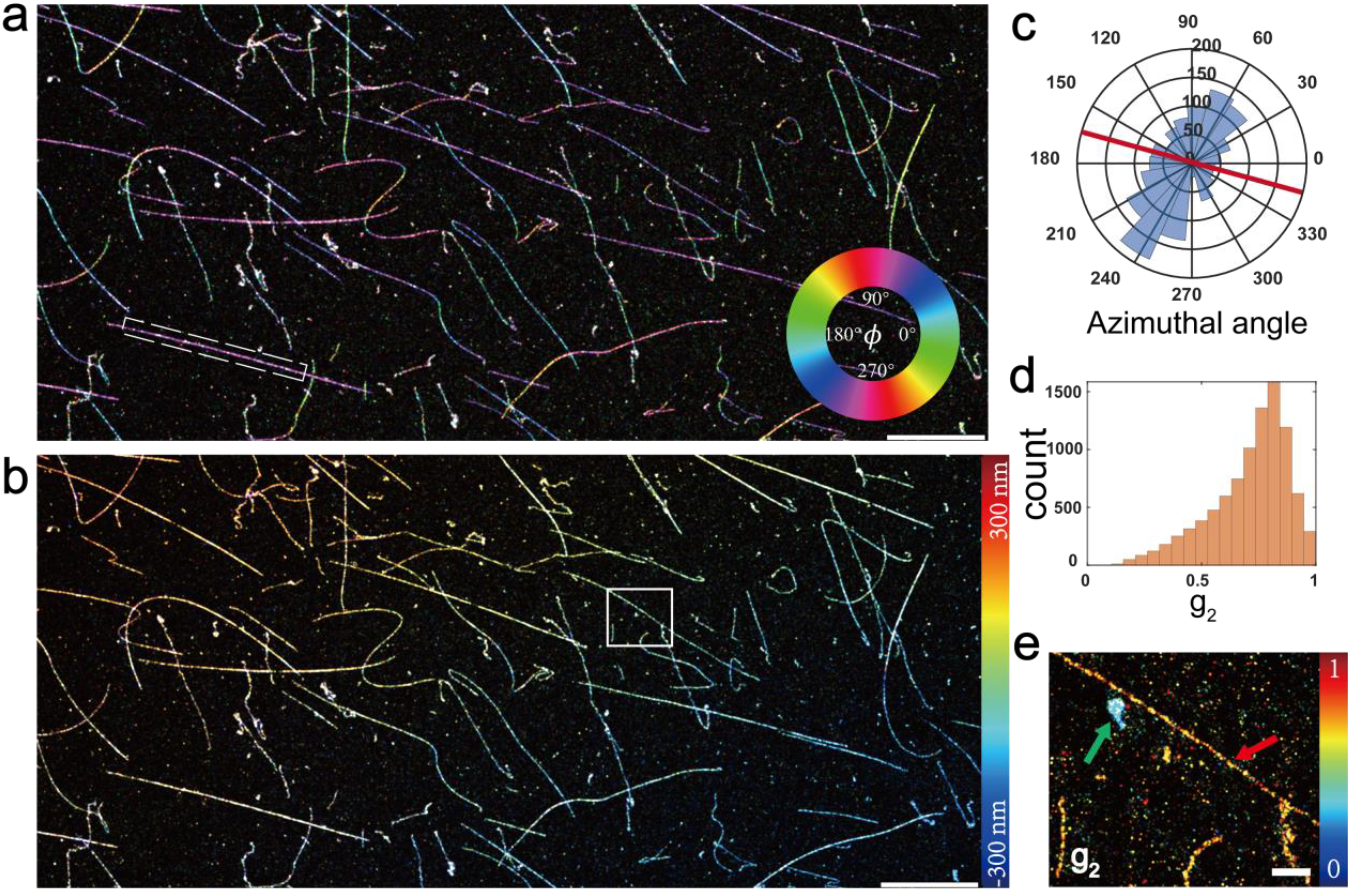
Simultaneous measurement of 3D positions and orientation information in λ-DNA using CBspline. **a**, Azimuthal angle images of Sytox Orange in λ-DNA. **b**, 3D super resolution images of the same region as in **a. c**, Azimuthal angle distribution in the dashed region in **a**. The red line represents the angle of inclination of the rectangular dashed line in **a. d**, *g*_2_ distribution of the same region as **c. e** *g*_2_ images of the boxed region in **b**. Scale bars, 10 μm (**a**,**b**), 1 μm (**e**).

## Discussion

In summary, we proposed a B-spline based localization algorithm for high-dimensional PSF and use this method to extract rich multi-dimensional single-molecule information. Thanks to the efficient memory storage scheme for B-spline interpolation, higher dimensional information of single molecules can be modeled and analyzed with spline interpolation. Both experimental measured and theoretical derived PSF models can be modeled with B-spline. The algorithm was also implemented in multiple channels framework with globLoc functionality which allows flexible parameters sharing between different channels to obtain maximum information from multiple channels data(*30*). Furthermore, we employed GPU to accelerate the algorithm and improved the analysis speed by 10∼100 times compared to CPU based algorithms.

Both in simulated and experimental data, we showed that B-spline based single molecule localization algorithms could precisely extract multi-dimensional information from single molecules and reach the theoretical minimum uncertainty, including 3D localizations, color information, interference phase information in 4Pi data and dipole orientation. We showed that our previously published DMO-PSF(*20*) could achieve both better 3D localization precision and color separation ability compared to that of the conventional astigmatism-based 3D PSF. Our IAB-based 4Pi-PSF model(*11*) can also be interpolated with B-splines without the need for developing specialized PSF fitters. Finally, we demonstrated the capability of B-spline based PSF models for simultaneous dipole orientation and 3D localization. All these results showed the great versatility of B-spline for modeling high dimensional PSFs with arbitrary shape.

Compared to the deep learning based single molecule analysis which needs a well-trained network for specialized PSF models and imaging conditions(*2, 32, 33*), our B-spline PSF fitter is ready to use for different PSF models and could return uncertainties for each parameter. The fitter is also capable of global fitting of multiple channel data (e.g., biplane, multicolor), allowing maximum information extraction across different channels. In addition, the GPU based B-spline fitter is open source and can be easily integrated in custom software. We anticipated that the fast and easy to use B-spline PSF fitter we developed here will enable complex PSF models with rich information content more accessible to the broad communities without the need for complicated PSF calculation.

## Materials and Methods

### Calculation of cubic B-spline interpolated PSFs

The nth order B-spline of N dimensional data are as follow:

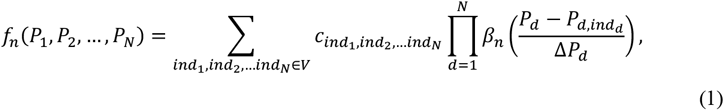

where *P*_1_, *P*_2_ … *P*_*N*_ is the parameters in each dimension (e.g., *x, y, z*); *ind*_*N*_ is the index of node in *N*th dimension. *V* is a set that contains the number of the nodes in all dimensions. 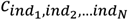 are the B-spline coefficients at node (*ind*_1_, *ind*_2_, … *ind*_*N*_). Δ*p*_*d*_ is the step size in *d* dimension (e.g., physical pixels size in x, y dimension); 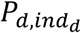 is the position of the *ind*_*d*_th node in dimension *d*. *β*_*n*_(*x*) denotes the nth order basis function which is given by:

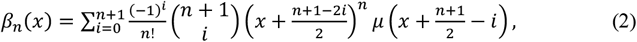

where

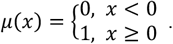

As confined by *μ*(*x*), only the four nearest basis functions for each node are non-zero for the cubic B-splines. To calculate the B-spline coefficients, we utilized a direct filter to extract them from the original discrete data(*17, 34*). The B-spline can be defined as a weighted summation of multiple basis functions at a sampled position. This constraint ensures a one-to-one correspondence between B-spline coefficients and exact interpolation. Consequently, the coefficients can be acquired by simple linear filtering(*34*).

In addition to computing the interpolation value, it is essential to calculate its partial derivative to obtain the optimization direction. An interesting characteristic of the B-spline is that its first-order partial derivatives can be expressed in a recursive form.

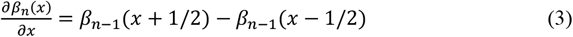

To compute the partial derivatives of *f* with respect to a specific dimension *d*, it is adequate to substitute the basis function in that dimension with its derivative. The derivative of a third-order basis function can be expressed as a polynomial of a second-order basis function. Its complete derivative formula is therefore expressed as

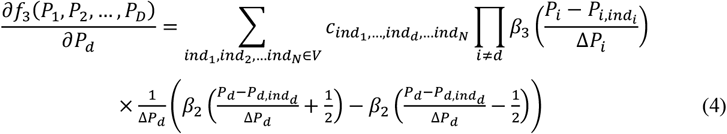

Since the above formula has a regular structure, it is quite straightforward to extend to higher dimensions. Besides, the number of coefficients is only related to number of the nodes. Therefore, the memory consumption of B-spline coefficients for higher dimensional PSFs does not significantly increase.

### Construction of 3D+λ PSF

To evaluate the performance of B-spline interpolated PSFs on the 4D PSF localization, we firstly constructed 3D+λ PSF both theoretically and experimentally. For the simulated 3D+λ PSF model, we employed a full vectorial PSF model(*35*) to generate both astigmatic and DMO PSF at different wavelengths (**Supplementary Note 3**). Data was synthesized with the following parameters: NA 1.5; pixel size 110 nm for x, y; refractive indices of the medium, cover glass, and immersion medium are 1.406, 1.525, and 1.518, respectively. PSFs at two emission wavelengths (580 nm and 680 nm) were investigated. Pupil function with different Zernike based aberrations was used to simulate the 3D PSFs. For astigmatism-based 3D PSF (**Supplementary Figs. 2, 4, 5 and 6b**), first order astigmatism 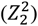 with 80 nm amplitude was employed. For the 1.2 µm DMO PSF (**Figs. 2b-f** and **Supplementary Fig. 6a**), Zernike aberrations with 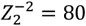 nm, 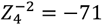 nm, *and* 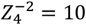 nm were employed. 3D PSFs with size of 31×31×41pixels, axial range of 400 nm above and below the focus (20 nm axial sampling step). The point emitter was place at 1µm away from the coverslip (z = 0 nm).

We then further padded the data in the fourth dimension by repeating the boundary elements to avoid discontinuity of the gradients. Before concatenating the two monochromatic 3D PSFs into a 4D PSF, we aligned these two PSFs using 3D cross-correlation to ensure a consistent coordinate between the PSF model of different color. To further reduce the effect of noise, we regularized the 4D PSF model by smoothing it in the z and color dimension with a smoothing B-Spline (smoothing coefficients lambda are 0.1 and 0.2, respectively)(*17*).

To further acquire the experimental 3D+λ PSF, we recorded the z stacks (ranging from - 400 nm to 400 nm with step of 20 nm) for different fluorescent color beads (40 nm diameter, red or dark red beads with emission maximum at 605 or 680 nm, F10720, Invitrogen) by illuminating with a 561 nm (or 640 nm) laser, individually. Here, the 3D DMO PSF was modulated by a deformable mirror (DM140A-35-P01, Boston Micromachines). For preparation of the bead samples, we firstly cleaned the precision coverslips (no. 1.5H, CG15XH, Thorlabs) by sequentially sonicating in 1 M potassium hydroxide (KOH), Milli-Q water and ethanol for 30 min, and then incubated the red (or dark red) beads on coverslips with 0.1 M MgCl_2_ for ∼5 min. To evaluate the color separation accuracy at different axial positions and photon emission experimentally (**Fig. 2d**), we moved the objective at different focal positions (from -400 nm to 400 nm with 200 nm step size) and gradually changed the illumination power at each z position during image acquisition.

### Construction of multi-channel 4Pi-PSF model

Four interference channel 4Pi PSF model was constructed in both theoretical simulation and experimental measurements. The theoretical simulation was conducted with a full vectorial PSF model (**Supplementary Note 3**) of both objectives using the following parameters: NA 1.35; refractive index 1.40 (immersion medium and sample) and 1.518 coverslip; emission wavelength 668 nm; pixel size 120 nm. Additional astigmatism 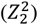 was added to the lower (100 mλ) and upper (-100 mλ) objectives separately. The sampling rate is 20 nm in the axial dimension and 0.1963 rad (equivalent to 10nm) in the phase dimension.

To construct an experimental four channel 4Pi-PSF model, 40 nm dimeter red fluorescence beads (F8793, Invitrogen) were imaged at z positions from −500 nm to 500 nm (20 nm intervals) with each z position recording 37 frames(*30*). We then calculated the I, A, B matrices (**Supplementary Note 4**) from these four interference channel calibration beads as described by our IAB-4Pi-PSF model(*11*). We then used these matrices to computer the complete 4D 4Pi-PSF at different interference phases (0.04π phase step between 0 to 2π). Considering that each channel has different aberrations, we built four 4Pi-PSF models with 19×19 ROI size and interpolate them with CBspline for each channel, separately.

### Construction of 5D vortex PSF model

Vortex PSF(*7*) was employed in this work to simultaneously estimate the 3D position and dipole orientations. As it is difficult to control the dipole orientation in a predefined manner, only theoretical PSF model was used. To simulate a theoretical vortex PSF, the following parameters were used: NA 1.47; the refractive index of immersion oil 1.518, coverslip 1.523, sample medium 1.33; emission wavelength 600 nm; pixel size 65 nm. Vortex phase-based pupil function was used to generate the 5D PSF with vectorial PSF theory (**Supplementary Note 3**). Here, the axial sampling rate of the vortex PSF is 20 nm between 500 nm above and below the focus in z direction, 10° step for azimuthal angle between 0° and 360°, and 2.5° step for polar angle between 0° and 90°. For the free rotating portion, we summed images of three orthogonal dipoles with equal strength to mimic the random orientation. The final PSF is the weighted sum of fixed dipole PSF and freely rotating PSF (**Supplementary Note 5**).

### GPU implementation of CBspline PSF fitter

We implemented the spline fitter with both Cspline and CBspline using CUDA C/C++ in NVIDIA CUDA® enable graphic cards. Each thread was pointed to a multi-dimensional single molecule data and calculated for each single molecule. We put the single molecule data and spline coefficients into the global memory of GPU and employed 64 threads in each block for the computation. Both the CPU-based C code and the GPU-based CUDA code were compiled with Microsoft Visual Studio 2019 into MEX files which could be called by MATLAB (MathWorks). For speed evaluation (**Supplementary Fig. 10, Fig. 1c**), we used an Intel Core i9-11900K CPU clocked at 3.5GHz for CPU fitting. An NVIDIA GeForce GTX 3080 graphics card with 10GB memory was used for GPU-based speed evaluation.

### Data process of dual-color NPC and mitochondria single molecule image

We’ve integrated a B-spline multi-dimensional fitter into the GUI we developed previously(*15*) (**Supplementary Fig. 12**). Here, we selected and cropped single molecule images with size of 15×15 by find the local maxima from the smoothed images, which filter by difference of gaussians. Segmented images were fitted by the CBspline fitter using the 3D+λ PSF model by the customized maximum likelihood estimation (**Supplementary Note 1**). For high accuracy of color classification, single molecules with photons less than 4,000 photons were discarded. Post processing steps like localization grouping for single molecules persisting over consecutive frames, drift correction and molecule filtering (rejection of molecules with low localization precision, low photons) were performed with SMAP(*36*).

### Sample preparation for dual-color imaging of NPC and mitochondria

U2OS cells (Nup96-SNAP, 300444, Cell Line Services) were grown in DMEM (10569, Gibco) containing 10% (v/v) fetal bovine serum (10270-106, Gibco), 1× MEM NEAA (11140-050, Gibco),100 U/ml penicillin and 100 μg/ml streptomycin (15140-122, Gibco). Cells were cultured in a humidified atmosphere with 5% CO_2_ at 37 °C and passaged every two or three days. For super-resolution imaging, cells were seeded on the cleaned coverslips which were further irradiated under ultraviolet light for 30 min and cultured for 2 d to achieve a confluency of ∼70%. Before labeling, cells were prefixed in 2.4% (w/v) paraformaldehyde (PFA) for 30 s, permeabilized in 0.4% (v/v) Triton X-100 for 3 min, further fixed in PFA for 30 min and quenched in 0.1 M NH_4_Cl for 5 min. We then sequentially labeled Nup96 proteins with SNAP ligand conjugated Alexa Fluor 647 (BG-AF647, S9136S, New England Biolabs) and mitochondria with primary anti-Tomm20 antibodies (rabbit polyclonal, ab78547, Abcam) and secondary antibodies conjugated CF568 (sab-CF568, SAB4600310, Sigma). For staining of Nup96, cells were blocked in Image-iT FX signal enhancer (I36933, Invitrogen) for 30 min, incubated in dye solution (1 μM BG-AF647, 1mM dithiothreitol and 0.5% (w/v) bovine serum albumin (BSA) in PBS) for 1 h, and washed 3 times in PBS for 5 min each. For staining of Tomm20, cells were incubated in PBS containing 1 μg/ml anti-Tom20 and 3% BSA overnight at 4 °C, washed 3 times in PBS for 5 min each, incubated in PBS containing 1 μg/ml sab-CF568 and 3% BSA for 1 h, and washed 3 times in PBS for 5 min each. Finally, cells were postfixed in 4% PFA for 10 min and stored in PBS at 4 °C before imaging.

### Microscopy

In this work, dual-color imaging was performed at room temperature (24°C) and by a custom-built microscope equipped with a deformable mirror(*20*). We used single-mode fiber (P3-405BPM-FC-2, Thorlabs) illumination for excitation. The fluorescence signal was imaged through a high numerical aperture oil immersion objective (NA 1.5, UPLAPO100XOHR, Olympus). Field-programmable gate array (Mojo, Embedded Micro) were used to control 561nm laser (MGL-FN-561nm, 300mW, CNI) and 640nm laser (iBEAM-SMART-640-S-HP, 200mW, TOPTICA Photonics) with microseconds pulsing accuracy. The fluorescence is separated from the excitation light by a dichroic mirror (ZT405/488/561/640rpcxt-UF2, Chroma) to avoid strong background effects and through two band pass filters (NF03-405/488/561/635E-25 and FF01-676/37-25, Semrock). After a 4f system consisting of two lenses (125 mm, 75 mm), it is reflected on a deformable mirror (DM140A-35-P01, Boston Micromachines) on the conjugate back focal plane, and then imaged with 110 nm pixel size on the camera (Dhyana 400BSIV3, Tucsen).

## Code availability

Source code for the software used in this manuscript is contained in **Supplementary Software 1** and updated versions can be freely downloaded at: https://github.com/Li-Lab-SUSTech/GPU-based-B-spline-Fitter

## Acknowledgements

The authors are grateful to Sjoerd Stallinga and Bernd Rieger from Delf University of Technology for providing us with the Vortex PSF based single molecule experimental data. This work was supported by the National Natural Science Foundation of China (62375116), Key Technology Research and Development Program of Shandong (2021CXGC010212), Shenzhen Science and Technology Innovation Commission (Grant No. JCYJ20220818100416036 and KQTD20200820113012029), Guangdong Natural Science Foundation Joint Fund (2020A1515110380), Guangdong Provincial Key Laboratory of Advanced Biomaterials (2022B1212010003), Startup grant from Southern University of Science and Technology.

## Author contributions

Y.L. conceived the concept and supervised the project. Y. L, M. L. and W. S. developed the methods and wrote the software. M. L., W. S., S. F., Y. F., and L. Z. acquired and analyzed the data. S. L. contributed to the 4Pi data analysis. Y. L., M. L. and W. S. wrote the manuscript.

## Data availability

All data are available upon reasonable request from the corresponding authors.

