## Supplementary file for "Fast and universal single molecule localization using multi-dimensional point spread functions"

| | |
|---|---|
| **Supplementary Figure 1** | Comparison of Cspline and CBspline interpolation on simulated data |
| **Supplementary Figure 2** | Comparison of accuracy between CBspline and Cspline in 3D localization |
| **Supplementary Figure 3** | Localization results for 3D position and interferometric phase of 4Pi-PSF |
| **Supplementary Figure 4** | Comparison of astigmatic PSFs at varying wavelengths |
| **Supplementary Figure 5** | Classification accuracy of simulated astigmatic PSF at different wavelengths |
| **Supplementary Figure 6** | Axial localization precision of DMO PSF and astigmatic PSF |
| **Supplementary Figure 7** | Photon Distributions of red and dark red beads in Figure 2d |
| **Supplementary Figure 8** | Photon distribution of the 3D imaging data in Figure 3 |
| **Supplementary Figure 9** | Schematic diagram of the 4Pi-PSF model |
| **Supplementary Figure 10** | Comparison of speed between CBspline and Cspline as a function of ROI size |
| **Supplementary Figure 11** | Localization precision of single molecule orientation, 3D position, and orientational constraint for Vortex PSF as function of dipole orientation ($\theta$ and $\phi$), z and g2, separately |
| **Supplementary Figure 12** | GUI of the multi-dimensional CBspline fitter |
| **Supplementary Table 1** | Comparison of memory needed for Cspline and CBspline based spline coefficients |
| **Supplementary Note 1** | Maximum likelihood estimation of multi-dimensional data |
| **Supplementary Note 2** | Calculation of multi-dimensional CRLB |
| **Supplementary Note 3** | Vectorial PSF model |
| **Supplementary Note 4** | 4Pi-PSF construction |
| **Supplementary Note 5** | Single molecule localization with orientation estimation |

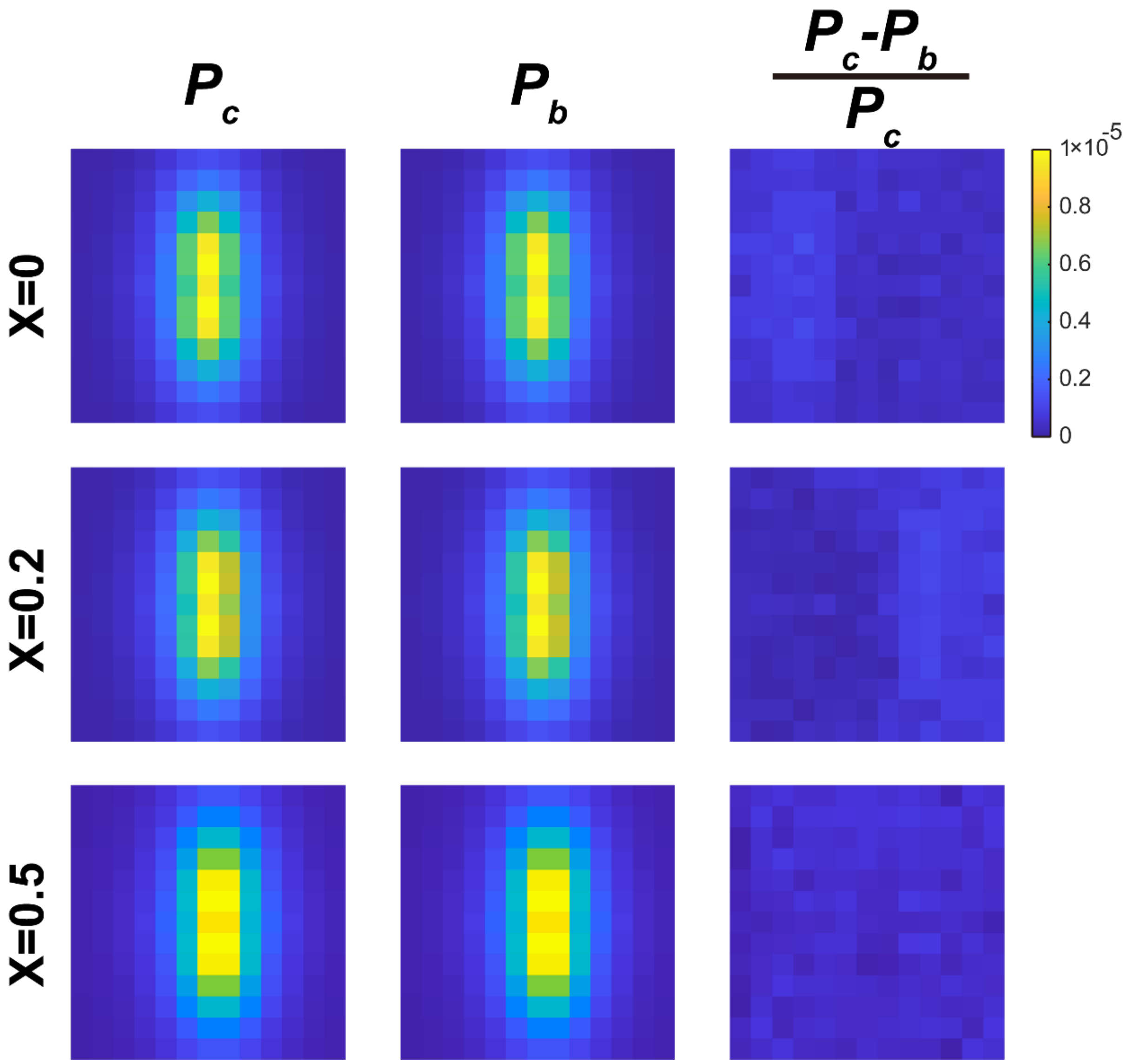


**Supplementary Figure 1: Comparison of Cspline and CBspline interpolation on simulated data.** The simulated astigmatic PSF was interpolated using the Cspline ($P_c$) and CBspline ($P_b$), respectively. The PSF images at the original position, as well as at lateral offset of 0.2 and 0.5 pixel, were interpolated using both methods, separately. Their relative differences are calculated.

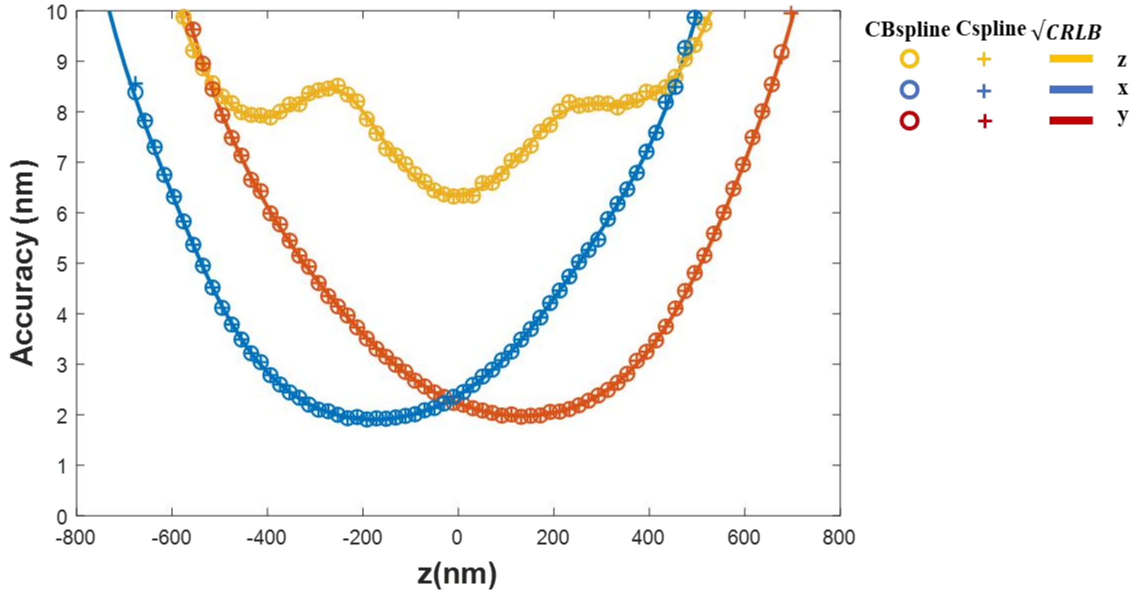


**Supplementary Figure 2: Comparison of accuracy between CBspline and Cspline in 3D localization.** Astigmatic PSF images were simulated using vectorial model with 5,000 photons and 20 background photons/pixel. Two fitting methods (based on Cspline and CBspline) were employed for 3D localization. Yellow, blue, and red represent x, y and z dimensions, respectively. The circle and plus sign represent the localization accuracy of the CBspline and Cspline fitters. Solid lines represent the CRLB of each dimension.

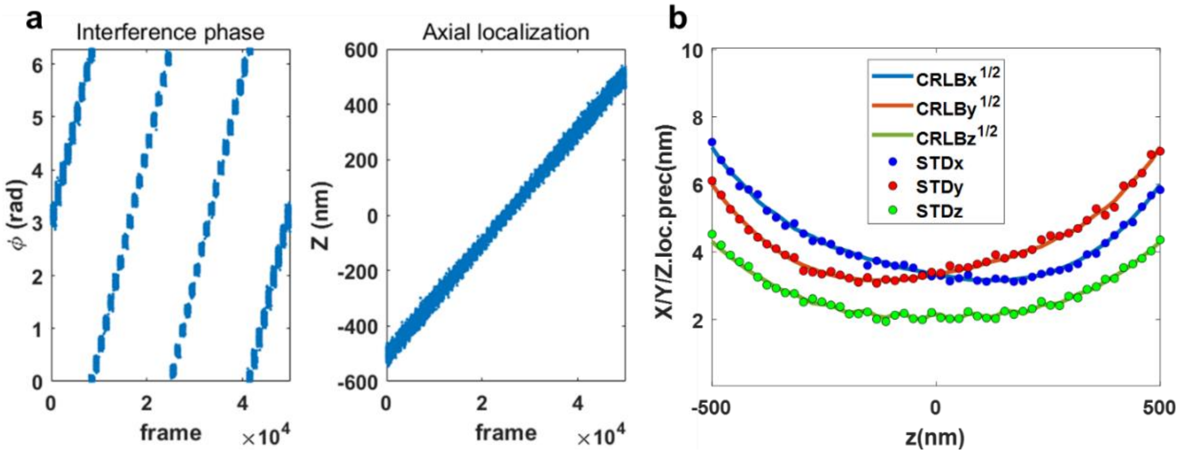


**Supplementary Figure 3: Localization results for 3D position and interferometric phase of 4Pi-PSF**. 5,000 4Pi single molecule images with 100 mλ astigmatism were simulated at axial range from -500 nm to 500 nm. Each objective collected 1,000 photons and 20 background photons/pixel. **a,** The localization results for the interference phase $\varphi$ (converted to radian units, left panel) and axial localization (right panel). **b,** CRLB and localization precision, where z dimension is calculated from the phase in **a**.

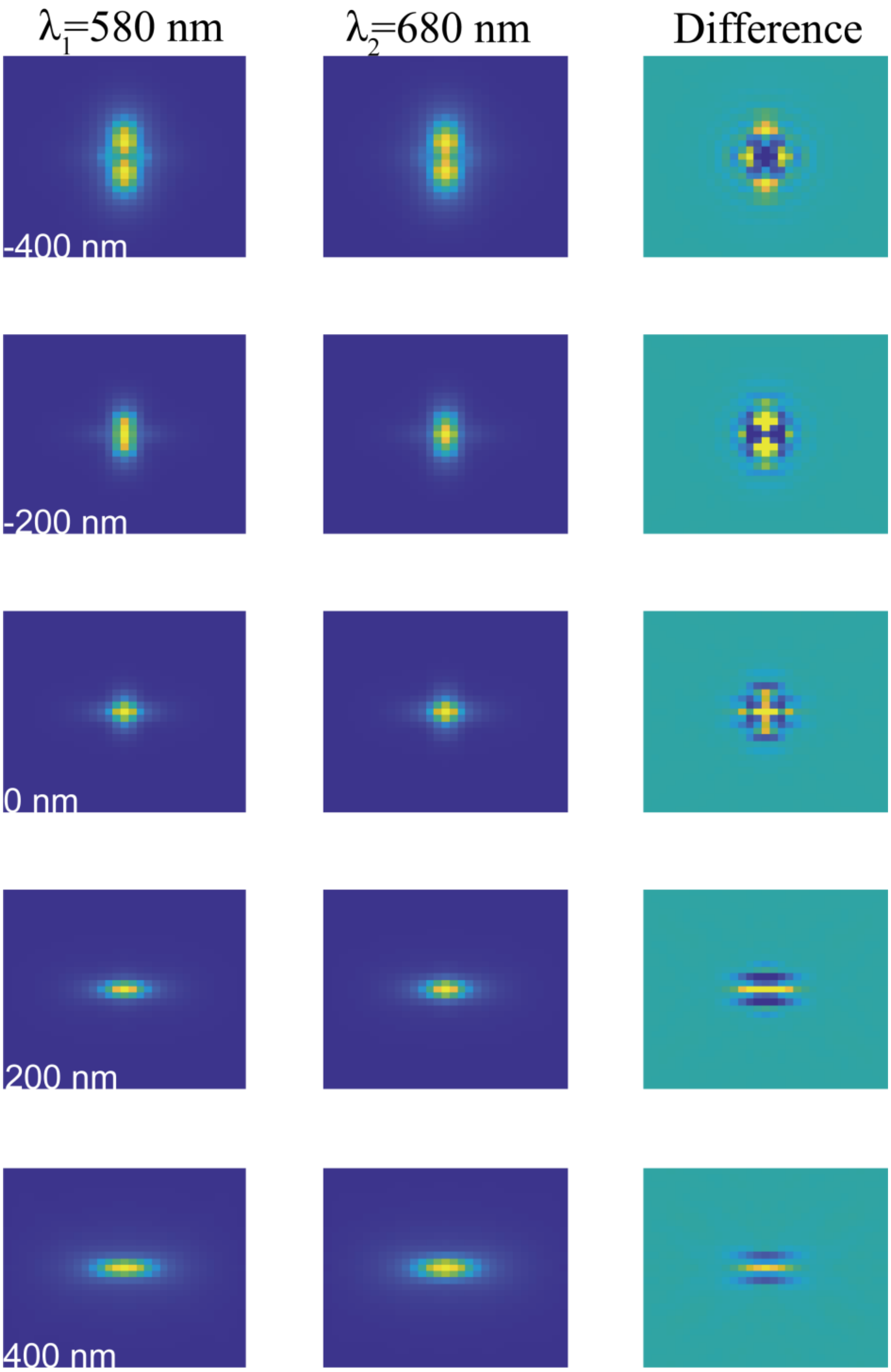


**Supplementary Figure 4: Comparison of astigmatic PSFs at varying wavelengths.** Astigmatic PSFs (80 nm astigmatism) with different emission wavelengths (580 nm and 680 nm) at different axial positions are generated using vectorial model. The difference is shown in the right column.

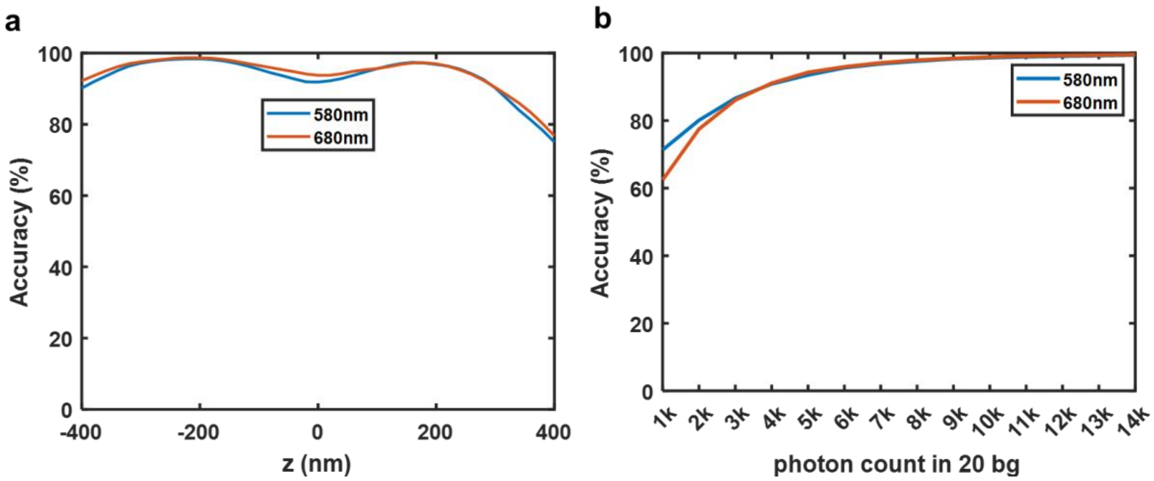


**Supplementary Figure 5: Classification accuracy of simulated astigmatic PSF at different wavelengths.** **a**, Color classification accuracy of single molecule images with 5,000 photons/localization and 20 background photons/pixel at different emission wavelengths (580 nm and 680 nm) within an axial position ranges from -400 nm to 400 nm. **b**, Average accuracy of color classification as a function of photon counts for single molecules in the z range 400 nm around focus.

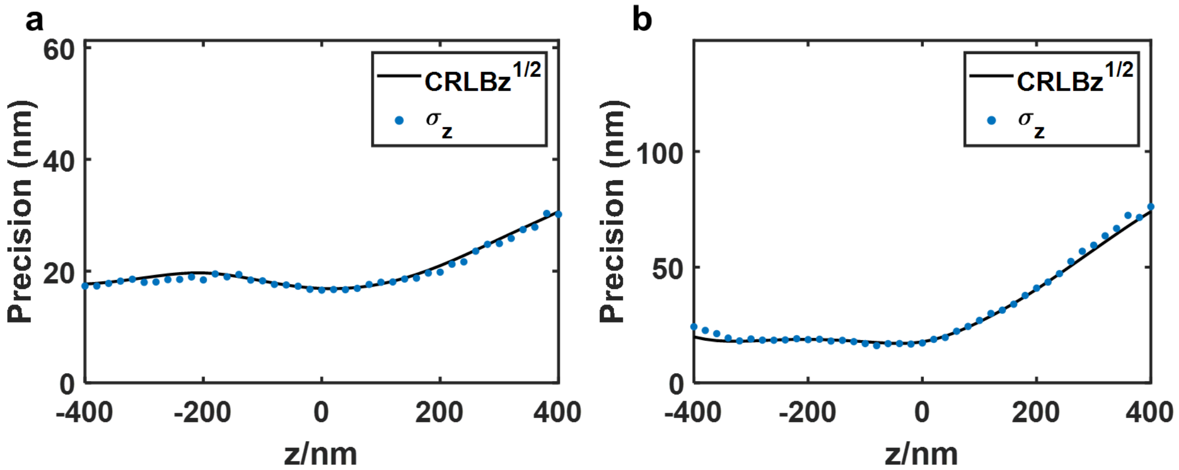


**Supplementary Figure 6: Axial localization precision of DMO PSF and astigmatic PSF. a,** z localization precision for DMO PSF. **b**, z localization precision for astigmatic PSF. Single molecules with 2,000 photons and 20 background photons/pixel were used for evaluation. Vector PSF model was employed. Here, the refractive indices for sample medium, cover glass and immersion oil are 1.406, 1.525 and 1.518, respectively. The single molecules (z = 0 nm) were located at 1 μm away from the coverslip.

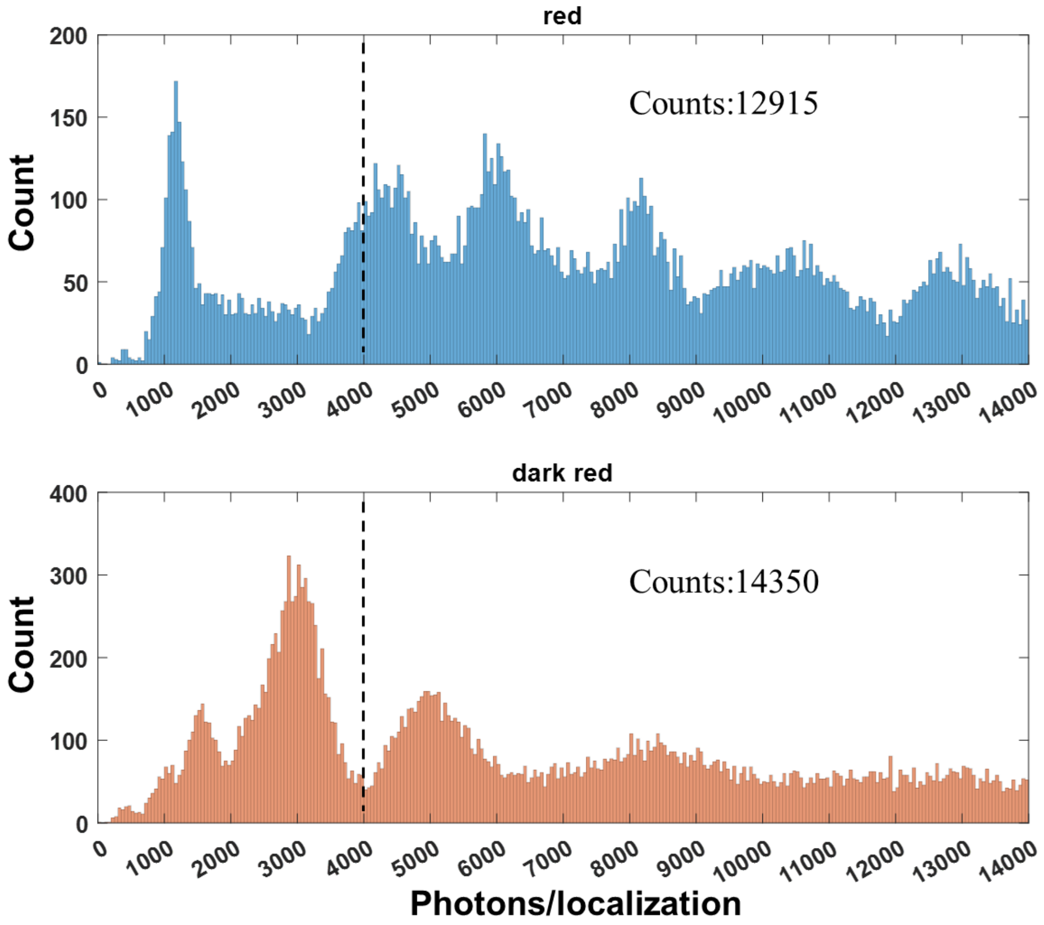


**Supplementary Figure 7: Photon Distributions of red and dark red beads in Figure 2d.** To evaluate the accuracy at different position, single bead with photon counts ranges from 4,000 to 14,000 were used to exclude the dim and aggregated beads. 12,915 red (605 nm) beads and 14,350 dark red (680 nm) beads were used for evaluation.

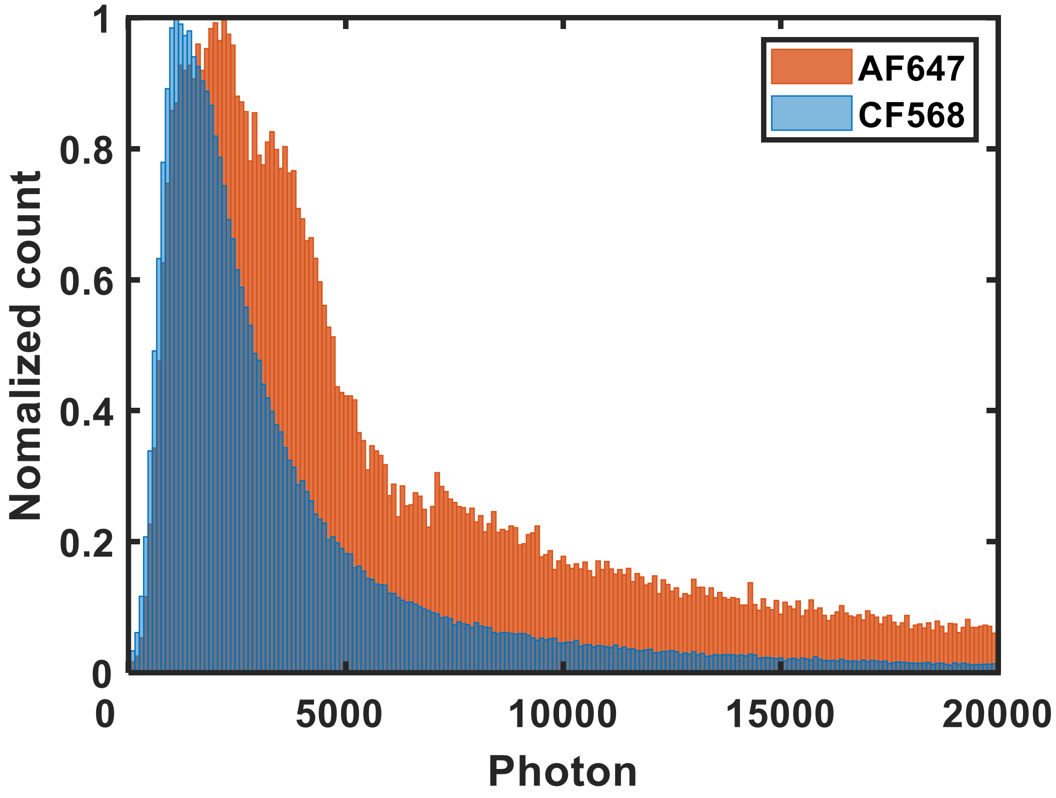


**Supplementary Figure 8: Photon distribution of the 3D imaging data in Figure 3.** The averaged photon count for the fluorescent dyes CF568 and AF647 are 5,264 and 9,393, respectively. The photon was calculated based on the grouped single molecule events.

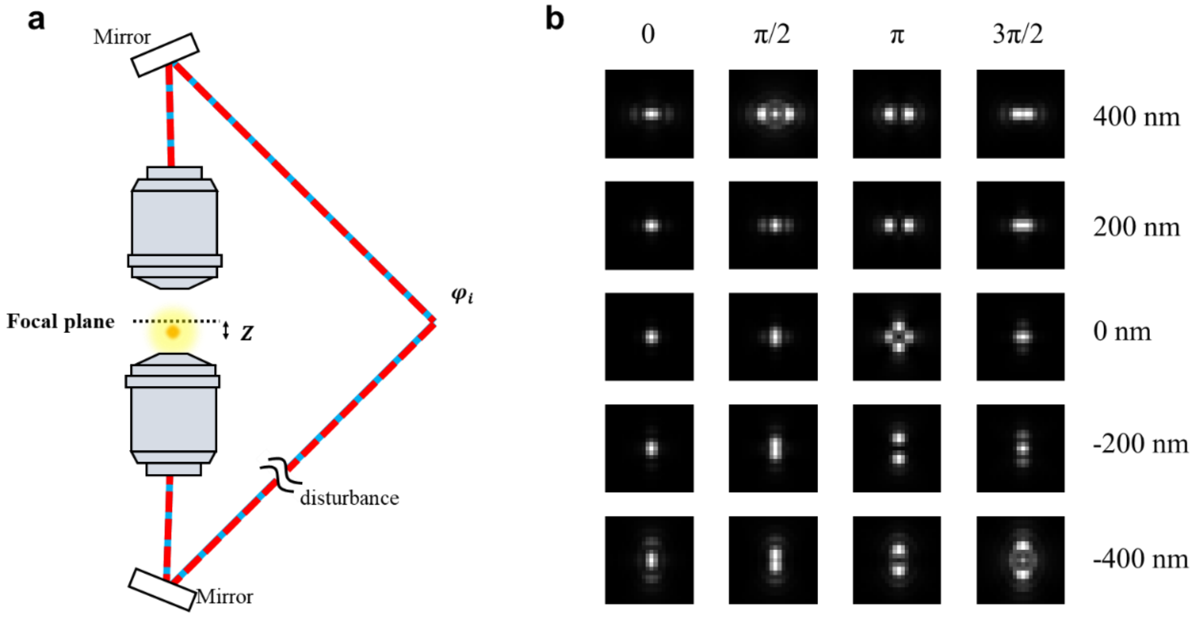


**Supplementary Figure 9: Schematic diagram of the 4Pi-PSF model. a**, Simplified illustration of the dual-objective 4Pi-SMLM system. **b**, 4Pi-PSFs with different interference phase (0, π/2, π, 3π/2) at different axial position.

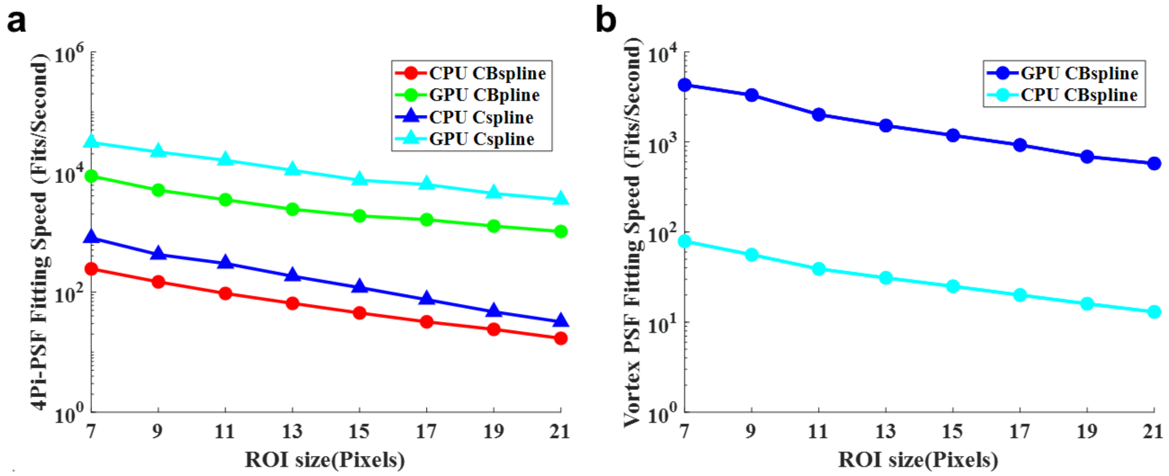


**Supplementary Figure 10: Comparison of speed between CBspline and Cspline as a function of ROI size. a,** Fitting speed of 4Pi-PSF with CBspline and Cspline. In the Cspline method, the IAB-4Pi-PSF model was used for 4D fitting as described before(*1*). **b**, Fitting speed of 5D Vortex-PSF with CBspline. Simulated single molecule images were generated using vector PSF model. Fits per second were measured on an i9-11900K CPU and a NVIDIA GeForce RTX 3080 consumer graphics card. The GPU code of CBspline is overall about 40 and 50 times faster than the CPU code of CBspline for 4Pi-PSF and Vortex PSF, respectively.

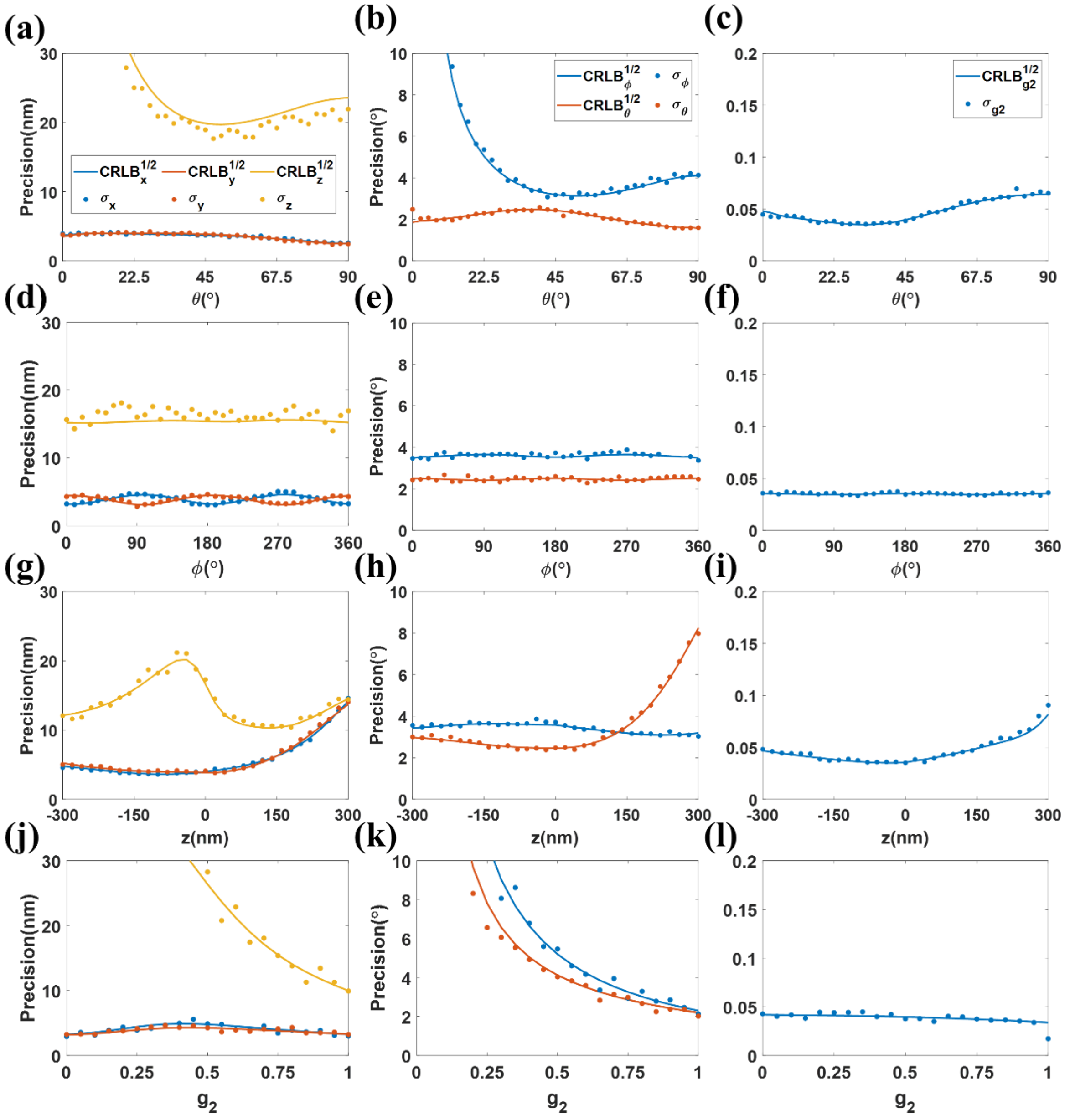


**Supplementary Figure 11: Localization precision of single molecule orientation, 3D position, and orientational constraint for Vortex PSF as function of dipole orientation (θ and ϕ), z and g₂, separately.** Vortex PSFs with 5,000 photons and 10 background photons/pixel were used for evaluation. Localization precision of x, y, z are shown in left column (**a**, **d**, **g**, **j**). Localization precision of dipole orientation are shown in the middle column (**b, e, h, k**). Localization precision of orientational constraint is shown in right column (**c, f, i, l**). PSFs of fixed dipole were used. The default values for the polar and azimuthal angels are 45°, z position is 0 nm, orientational constraint g2 is 0.75 if they were not specified in the figure.

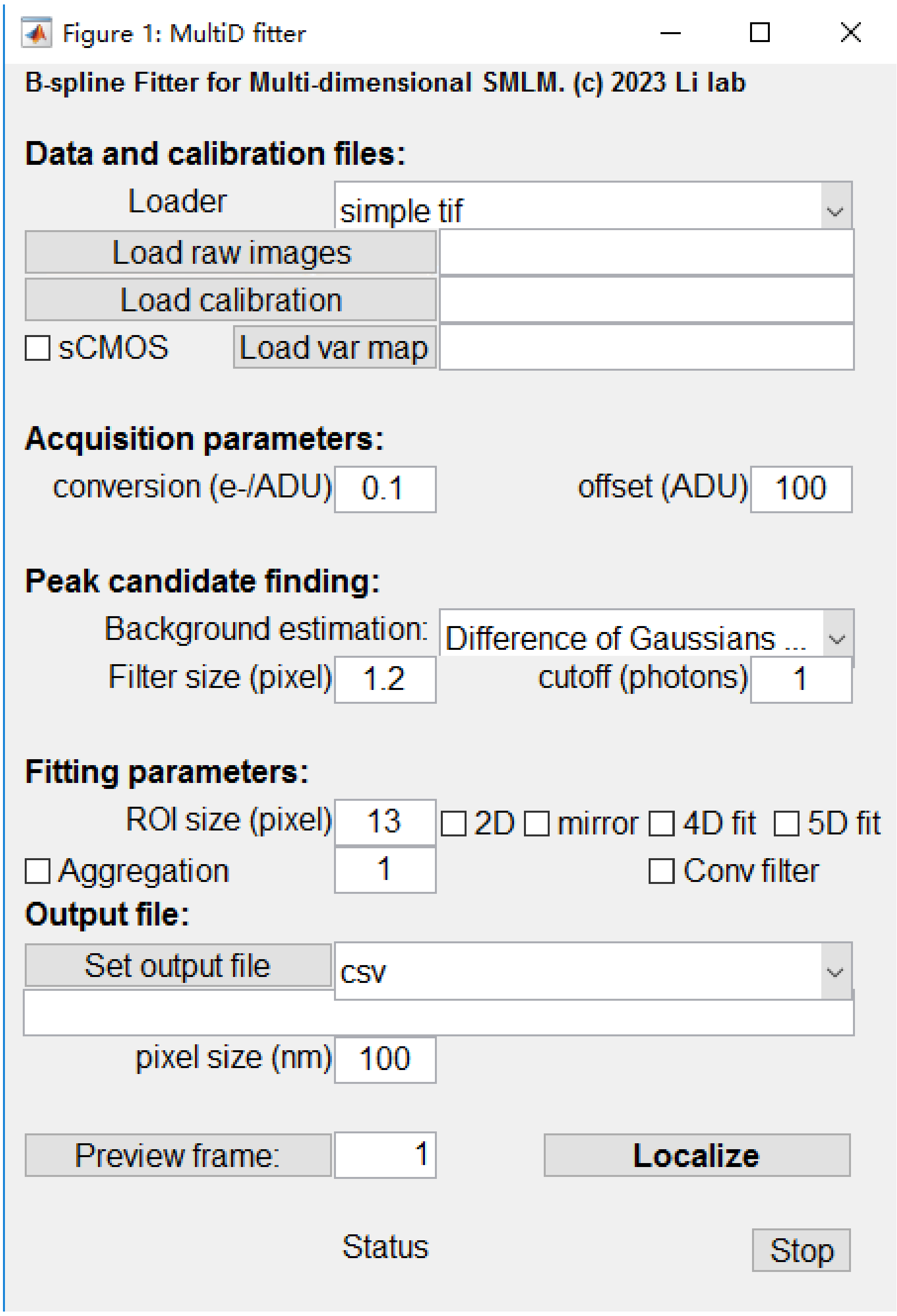


**Supplementary Figure 12: GUI of the multi-dimensional CBspline fitter.** We have incorporated high dimensional localization capabilities into the 3D Cspline single molecule localization fitter(*2*). After adding the raw image and calibration coefficients, the high dimensional localization can be selected by ticking either the 4D fit or the 5D fit. The preview frame button can also be used to preview the result of each frame before finalizing the localization by Localize.

**The size of memory required**

| Method | 3D data (31*31*50) | 4D data (31*31*50*50) | 5D data (31*31*50*50*50) |
|---|---|---|---|
| CBspline | 239.3kB | 12.3MB | 656.5MB |
| Cspline | 11.7MB | 2.3GB | 458.2GB |

**Supplementary Table 1: Comparison of memory needed for Cspline and CBspline based spline coefficients.** The memory of coefficients required for CBspline and Cspline. For CBspline, the data was further padded by replication in each dimension before calculating the spline coefficients. Float data type was used.

**Supplementary Note 1: Maximum likelihood estimation of multi-dimensional data**

We customized a maximum likelihood estimation (MLE) algorithm based on B-spline PSF model to flexibly localize multidimensional single-molecule data. The objective function of MLE across different dimensions is given by:

$$\chi^2 = 2\left(\sum_k(\mu_k - M_k) - \sum_k M_k * \ln\left(\frac{\mu_k}{M_k}\right)\right). \tag{s1}$$

Here, $M_k$ and $\mu_k$ are the measured and expected photon number on *kth* pixel, respectively. By minimizing $\chi^2$, we can obtain the maximum likelihood for the Poisson process. Nonlinear optimization methods typically involve iterative processes. In this case, we utilized a modified Levenberg-Marquardt (L-M) algorithm to optimize the parameters through iteration, as has been done previously(*2-4*):

$$(H_{i,j} + \lambda I)\Delta\theta_i = J_i \tag{s2}$$

$\Delta\theta_i$ is the update applied to the parameter indexing x, y, z, photons, background and extended estimated dimension, $\lambda$ is the damping factor and $I$ is a diagonal matrix equal to the diagonal elements of the Hessian matrix. $H_{i,j}$ is the Hessian matrix ignoring the second derivatives, defined as:

$$H_{i,j} = \sum_k \frac{\partial\mu_k}{\partial\theta_i}\frac{\partial\mu_k}{\partial\theta_j}\frac{M_k}{\mu_k^2} \tag{s3}$$

$J_i$ is the Jacobian matrix, defined as:

$$J_i = \sum_k \frac{\partial\mu_k}{\partial\theta_i}\frac{M_k-\mu_k}{\mu_k} \tag{s4}$$

where $\theta_i$ corresponds to the *ith* fitted parameter, which could pertain to molecule positions, background, photon count, or other relevant functional parameters. For multi-channel 4Pi-single molecule data with global shared parameters(*5*), the Hessian matrix and Jacobian matrix can be extended to:

$$H_{i,j} = \begin{cases} \sum_c\sum_k \frac{\partial\mu_{k,c}}{\partial\theta_{i,c}}\frac{\partial\mu_{k,c}}{\partial\theta_{j,c}}\frac{M_{k,c}}{\mu_{k,c}^2}, (\theta_{i,c}, \theta_{j,c} \in \boldsymbol{\theta_p}) \\ \sum_k \frac{\partial\mu_{k,c}}{\partial\theta_{i,c}}\frac{\partial\mu_{k,c}}{\partial\theta_{j,c}}\frac{M_{k,c}}{\mu_{k,c}^2}, (\theta_{i,c} \in \boldsymbol{\theta_p},\ \theta_{i,c} \in \boldsymbol{\theta_{qi}}) \\ \sum_k \frac{\partial\mu_{k,c}}{\partial\theta_{i,c}}\frac{\partial\mu_{k,c}}{\partial\theta_{j,c}}\frac{M_{k,c}}{\mu_{k,c}^2}, (\theta_{i,c} \in \boldsymbol{\theta_{qi}},\ \theta_{i,c} \in \boldsymbol{\theta_{qi}}, i = j) \\ 0, (\theta_{i,c} \in \boldsymbol{\theta_{qi}},\ \theta_{i,c} \in \boldsymbol{\theta_{qi}}, i \neq j) \end{cases}. \tag{s5}$$

$$J_i = \begin{cases} \sum_c\sum_k \frac{\partial\mu_{k,c}}{\partial\theta_{i,c}}\frac{(M_{k,c}-\mu_{k,c})}{\mu_{k,c}}, (\theta_{i,c} \in \boldsymbol{\theta_p}) \\ \sum_k \frac{\partial\mu_{k,c}}{\partial\theta_{i,c}}\frac{(M_{k,c}-\mu_{k,c})}{\mu_{k,c}} (\theta_{i,c} \in \boldsymbol{\theta_{qi}}) \end{cases}. \tag{s6}$$

where $\theta_{i,c}$ is the parameter $i$ in channel $c$, $M_{k,c}$ and $\mu_{k,c}$ are the measured and expected photon number on *kth* pixel in channel $c$. $\boldsymbol{\theta_p}$ and $\boldsymbol{\theta_{qi}}$ are the set of shared and non-shared parameters, respectively. Here, the scaling factor of each parameter between different channels is set as 1 by default.

### Supplementary Note 2: Calculation of multi-dimensional CRLB

To quantify the localization precision of the multi-dimensional B-spline fitter, the Cramér-Rao lower bound (CRLB) was utilized to indicate the lowest variance that our B-spline estimator can achieve. The CRLB can be calculated using the inverse of the Fisher information matrix:

$$var(\hat{\theta}_i) \geq CRLB_{\theta_i} = [I(\boldsymbol{\theta})^{-1}]_{ii}. \tag{s7}$$

Here $\hat{\theta}_i$ represents the estimation for the parameter $\theta_i$ of interest. The Fisher information matrix is defined as:

$$I_{i,j} = \sum_k \frac{1}{\mu_k} \frac{\partial \mu_k}{\partial \theta_i} \frac{\partial \mu_k}{\partial \theta_j}, \tag{s8}$$

For multi-channel fitting, the globLoc(5) method was utilized to fit shared parameters, resulting in the Fisher matrix being as follows:

$$I_{i,j} = E\left[\sum_c \sum_k \frac{\partial \chi^2}{\partial \theta_{i,c}} \frac{\partial \chi^2}{\partial \theta_{j,c}}\right] = \begin{cases} \sum_c \sum_k \frac{1}{\mu_{k,c}} \frac{\partial \mu_{k,c}}{\partial \theta_{i,c}} \frac{\partial \mu_{k,c}}{\partial \theta_{j,c}}, (\theta_{i,c}, \theta_{j,c} \in \boldsymbol{\theta_p}) \\ \sum_k \frac{1}{\mu_{k,c}} \frac{\partial \mu_{k,c}}{\partial \theta_{i,c}} \frac{\partial \mu_{k,c}}{\partial \theta_{j,c}}, (\theta_{i,c} \in \boldsymbol{\theta_p},\ \theta_{i,c} \in \boldsymbol{\theta_{qi}}) \\ \sum_k \frac{1}{\mu_{k,c}} \frac{\partial \mu_{k,c}}{\partial \theta_{i,c}} \frac{\partial \mu_{k,c}}{\partial \theta_{j,c}}, (\theta_{i,c} \in \boldsymbol{\theta_{qi}},\ \theta_{i,c} \in \boldsymbol{\theta_{qi}}, i = j) \\ 0, (\theta_{i,c} \in \boldsymbol{\theta_{qi}},\ \theta_{i,c} \in \boldsymbol{\theta_{qi}}, i \neq j) \end{cases}. \tag{s9}$$

**Supplementary Note 3: Vectorial PSF model**

To accurately model the PSF of the microscope with a high numerical aperture (NA) objective, we utilized a vectorial PSF model that incorporates multiple medium interfaces. The PSF of a fixed dipole is calculated by summing images of three orthogonal electrical field, which can be expressed as follows:

$$PSF \propto \sum_{l=x,y}\left|\sum_{d=x,y,z}\mu_d\ \mathcal{F}_{2D}\left\{A(\vec{\rho})e^{i\psi_{psf}}e^{i\psi_{pos}}E_{l,d}^{pupil}\right\}\right|^2 \quad \text{(s10a)}$$

$$\mu = \left[sin\theta_p cos\varphi_p,\ sin\theta_p sin\varphi_p,\ cos\theta_p\right], \quad \text{(s10b)}$$

where $\mathscr{F}_{2D}$ denotes the 2D Fourier transform. $\mu$ is the transition dipole moment which projects the dipole onto Cartesian unit vectors. $\varphi_p$ is the azimuthal angle and $\theta_p$ is polar angle of the dipole orientation. Considering that fluorescent probes are typically attached to the sample of interest in a flexible manner and can freely rotate, we can treat them as uncorrelated dipoles. Consequently, $\mu$ is 1/3 for a freely rotating emitter. $A(\vec{\rho}) = (1 - |\vec{\rho}|^2 NA^2/ {n_{med}}^2)^{-1/4}$ is an amplitude function considering aplanatic correction factor. $(\rho, \varphi)$ is the normalized polar coordinate in the pupil plane with $|\vec{\rho}| = 1$ corresponding to the max aperture angle $NA/n_{imm}$. $n_{med}$, $n_{cov}$ and $n_{imm}$ is the refractive index of sample medium, cover slip and immersion oil, respectively. $\psi_{psf}$ is the modulated phase for PSF engineering and $\psi_{pos}$ is the phase shift determined by emitter position:

$$\psi_{pos} = \frac{2\pi}{\lambda}\left(NAx_0\vec{\rho} * \hat{x} + NAy_0\vec{\rho} * \hat{y} + n_{med}z_0\sqrt{1 - (\frac{|\vec{\rho}|NA}{n_{med}})^2} - n_{imm}z_{depth}\sqrt{1 - (\frac{|\vec{\rho}|NA}{n_{imm}})^2}\right) \quad \text{(s11)}$$

where $\lambda$ is the wavelength, $(x_0, y_0, z_0)$ is the emitter position. $\hat{x}$ and $\hat{y}$ are the unit vectors of the x and y directions in pupil plane.$z_0$ represents the distance between the emitter and the cover glass. $z_{depth}$ is the distance between the nominal focal plane and the cover glass. $E_{l,d}^{pupil}$ represents the polarization vector with components $l = x, y$ in the pupil plane contributed by each dipole components $d = x, y, z$ separately:

$$E_{x,d}^{pupil} = t_p\vec{P}_d cos\varphi - t_s\vec{S}_d sin\varphi \quad \text{(s12)}$$

$$E_{y,d}^{pupil} = t_p\vec{P}_d sin\varphi + t_s\vec{S}_d cos\varphi \quad \text{(s13)}$$

$$with\left(\vec{P}=\begin{bmatrix}cos\,\theta_{med}\,cos\,\varphi\\cos\,\theta_{med}\,sin\,\varphi\\-\,sin\,\theta_{med}\end{bmatrix},\vec{S}=\begin{bmatrix}-\,sin\,\varphi\\cos\,\varphi\\0\end{bmatrix}\right) \quad \text{(s14)}$$

where $\vec{P}$ and $\vec{S}$ are the unit polarization vectors. $t_p$ and $t_s$ are the total Fresnel transmission coefficients for the p- and s-polarized light through multiple mediums: $t_{k=p,s} = t_{k,med-cov} \times t_{k,cov-imm}$. $\theta$ and $\varphi$ are the polar and azimuthal angles of the wave direction. Take the boundary $med - cov$ as an example, the Fresnel transmission coefficients are given by:

$$t_{p,\mathrm{med-cov}} = \frac{2n_{\mathrm{med}}\,cos\,\theta_{\mathrm{med}}}{n_{\mathrm{med}}\,cos\,\theta_{\mathrm{cov}}+n_{\mathrm{cov}}\,cos\,\theta_{\mathrm{med}}} \quad \text{(s15a)}$$

$$t_{s,\mathrm{med-cov}} = \frac{2n_{\mathrm{med}}\,cos\,\theta_{\mathrm{med}}}{n_{\mathrm{med}}\,cos\,\theta_{\mathrm{med}}+n_{\mathrm{cov}}\,cos\,\theta_{\mathrm{cov}}} \quad \text{(s15b)}$$

Finally, chirp *z*-transform was used to perform the 2D Fourier transform which allows breaking the relationship of the sampling points between imaging space and Fourier space.

**Supplementary Note 4: 4Pi-PSF construction**

To construct the 4Pi-PSF model, we employed the IAB model proposed in our previous work(*1*):

$$PSF_{4Pi}(\boldsymbol{r},\varphi) = I(\boldsymbol{r}) + A(\boldsymbol{r})cos(\varphi) + B(\boldsymbol{r})\sin(\varphi), \quad (s16)$$

where $\boldsymbol{r}$ is 3D coordinates (x, y, z) of emitter, while $\varphi$ denotes the phase difference that depends on the optical path difference between the two objectives. As described in our previous work(*1*), the 4Pi-PSF, which is derived from two single objective PSFs with different phases, can be easily decoupled into three matrixes (*I*, *A*, *B*). To modulate the interference phase of this four-channel PSF, we added a linear phase change from 0-2π to each channel.

**Supplementary Note 5: Single molecule localization with orientation estimation**

To enable the B-spline fitter in 3D localization microscopy with orientation awareness, another three parameters should be fitted: the polar angle, azimuthal angle, and degree of constraint. According to the previous method(*6*), we used a weighted summation of the freely rotating PSF $P_{free}$ and the orientation-fixed PSF $P_{fixed}$ to describe the final image formation model:

$$PSF_{vortex}(\boldsymbol{r},\theta,\phi,g_2,N,b)=\frac{N}{3}[(1-g_2)P_{free}(\boldsymbol{r})+g_2\,P_{fixed}(\boldsymbol{r},\theta,\phi)]+b. \quad \text{(s17)}$$

Here, $\boldsymbol{r}$ is the three-dimensional spatial position of the single molecule, $\theta$ and $\phi$ represent the orientation of the fixed dipole, $N$ represents the number of photons, and $b$ represents the background signal received at each pixel. The parameter g2 represents the degree of rotational constraint. g2 with a value closer to 1 indicates that the physical behavior is more like a fixed dipole, while a value closer to 0 indicates behavior more similar to a freely rotating dipole. The partial derivatives of the imaging model with respect to each parameter are given as follows:

$$\frac{\partial P_{vortex}}{\partial Ori}=\frac{N*g_2}{3}*\frac{\partial P_{fixed}}{\partial Ori}\ , \quad \text{(s18)}$$

$$\frac{\partial P_{vortex}}{\partial r}=\frac{N}{3}*[g_2\frac{\partial P_{fixed}}{\partial r}+(1-g_2)\frac{\partial P_{free}}{\partial r}]\ , \quad \text{(s19)}$$

$$\frac{\partial P_{vortex}}{\partial N}=\frac{1}{3}g_2P_{fixed}+\frac{1}{3}(1-g_2)P_{free}\ , \quad \text{(s20)}$$

$$\frac{\partial P_{vortex}}{\partial b}=1\ , \quad \text{(s21)}$$

$$\frac{\partial P_{vortex}}{\partial g_2}=\frac{1}{3}\left(P_{fixed}-P_{free}\right)\ . \quad \text{(s22)}$$
